# Pathways that affect anterior morphogenesis in *C. elegans* embryos

**DOI:** 10.1101/2023.04.23.537986

**Authors:** Balasubramaniam Boopathi, Irini Topalidou, Melissa Kelley, Sarina M. Meadows, Owen Funk, Michael Ailion, David S. Fay

## Abstract

During embryogenesis the nascent *Caenorhabditis elegans* epidermis secretes an apical extracellular matrix (aECM) that serves as an external stabilizer, preventing deformation of the epidermis by mechanical forces exerted during morphogenesis. We showed that two conserved proteins linked to this process, SYM-3/FAM102A and SYM-4/WDR44, colocalize to intracellular and membrane-associated puncta and likely function together in a complex. Proteomics data also suggested potential roles for FAM102A and WDR44 family proteins in intracellular trafficking, consistent with their localization patterns. Nonetheless, we found no evidence to support a clear function for SYM-3 or SYM-4 in the apical deposition of two aECM components, FBN-1 and NOAH. Surprisingly, loss of MEC-8/RBPMS2, a conserved splicing factor and regulator of *fbn-1*, had little effect on the abundance or deposition of FBN-1 to the aECM. Using a focused screening approach, we identified 32 additional proteins that likely contribute to the structure and function of the embryonic aECM. Lastly, we examined morphogenesis defects in embryos lacking *mir-51* microRNA family members, which display a related embryonic phenotype to *mec-8; sym* double mutants. Collectively, our findings add to our knowledge of pathways controlling embryonic morphogenesis.

**SUMMARY STATEMENT:** We identify new proteins in apical ECM biology in *C. elegans* and provide evidence that SYM-3/FAM102A and SYM-4/WDR44 function together in trafficking but do not regulate apical ECM protein deposition.

## INTRODUCTION

The development of embryos requires the tight coordination of cell proliferation, cell differentiation, cell movements, and tissue morphogenesis. Whereas many of the genes and mechanisms underlying proliferation, differentiation, and migration have been characterized in considerable detail, the morphogenesis of tissues, organs, and whole organisms is less well understood. This is likely due to the complexity of the morphogenetic process, constraints associated with genetic approaches including genetic redundancy, challenges with carrying out in vivo imaging and manipulations, and technical hurdles or limitations inherent to ex vivo studies.

The nematode *Caenorhabditis elegans* is a useful model in which to study various aspects of development including the morphogenesis of embryos and organs. Whereas certain aspects of *C. elegans* embryonic morphogenesis—such as gastrulation (Goldstein and Nance, 2020), embryonic elongation (Vuong-Brender et al., 2016), and dorsoventral enclosure (Chisholm and Hardin, 2005; Chisholm and Hsiao, 2012)—have been reasonably well described, the development and morphogenesis of the anterior embryo, in particular the epidermis and its integration with the oral cavity, have just begun to be characterized in detail (Grimbert et al., 2021; Portereiko and Mango, 2001).

Work from our lab and others has shown that *C. elegans* SYM-3, SYM-4, and MEC-8 act in a coordinated manner to prevent deformation of the anterior epidermis and internalization of the buccal cavity (mouth) in response to pharyngeal pulling forces during embryogenesis (Davies et al., 1999; Kelley et al., 2015; Yochem et al., 2004). Specifically, *mec-8; sym-3* and *mec-8; sym-*4 double mutants display a <u>p</u>harynx <u>in</u>gressed (Pin) phenotype and consequent larval lethality due to an inability to feed. In contrast, *mec-8*, *sym-3*, and *sym-4* single mutants are largely viable, as are *sym-3 sym-4* double mutants. These genetic results suggest that MEC-8 may act in a pathway or process that functions in parallel to the SYMs, whereas SYM-3 and SYM-4 may function together in a common pathway or complex.

Additional findings indicate that MEC-8, an ortholog of the mammalian RBPMS2 slicing factor (Lundquist et al., 1996), controls the alternative splicing of multiple RNA targets including *fbn-1* (Kelley et al., 2015; Spike et al., 2002), a protein related to vertebrate fibrillins (Frand et al., 2005). Moreover, misregulation of *fbn-1* is a key factor in causing the Pin phenotype of *mec-8; sym* double mutants (Kelley et al., 2015). FBN-1 functions as a structural component of the embryonic sheath—a specialized embryonic pre-cuticle—and in the larval cuticle, in which it is required for normal molting (Cohen et al., 2020b; Frand et al., 2005; Katz et al., 2022; Kelley et al., 2015). Both the sheath and cuticle are a category of apical extracellular matrix (aECM) that is secreted in large part by the epidermis (Page and Johnstone, 2007). Whereas the larval and adult cuticles serve as an exoskeleton and protect worms from the environment (Johnstone, 1994; Page and Johnstone, 2007), the sheath functions as a semi-rigid scaffold to prevent the embryonic epidermis from deformation by mechanical forces, including inward-pulling forces exerted by the elongating pharynx (Kelley et al., 2015; Priess and Hirsh, 1986; Vuong-Brender et al., 2017). Abnormal pharyngeal ingression has also been observed in *dex-1* and *dyf-7* mutants (Cohen Sundaram, 2019), which, like *fbn-1*, encode components of the aECM (Heiman and Shaham, 2009). Moreover, additional roles for the aECM in various aspects of morphogenesis and cell shape control have come to light, emphasizing the importance of this relatively understudied extracellular component of developing tissues (Cohen and Sundaram, 2020; Labouesse, 2012; Lamkin and Heiman, 2017; Li Zheng et al., 2020; Sundaram and Cohen, 2017).

Whereas the roles of MEC-8 family members in splicing are well established, the molecular and cellular functions of SYM-3 and SYM-4 are less known. SYM-4 is a predicted β-propeller protein with seven WD-repeats, suggesting a role in coordinating protein interactions. The mammalian ortholog of SYM-4, WDR44/RAB11BP/Rabphilin-11, was identified independently by two groups as a binding partner and candidate effector of RAB11 (Mammoto et al., 1999; Zeng et al., 1999), a highly conserved GTPase involved in several aspects of cell trafficking and cell biology (Ferro et al., 2021; Guichard et al., 2014; Welz et al., 2014; Zhang et al., 2021). Although more-recent studies have expanded these initial findings (Chung et al., 2014; Lucken-Ardjomande Hasler et al., 2020; Thibodeau et al., 2023; Walia et al., 2019), the full range of WDR44 functions largely remains to be elucidated. Even less is known about SYM-3, the ortholog of FAM102A/EEIG1. FAM102A family proteins contain an N-terminal C2 (NT-C2) domain, a motif involved in lipid binding that is found in membrane-associated proteins including the conserved trafficking factors EHBP1 and Synaptotagmin-1 (Lynch et al., 2007; Shi et al., 2010; Zhang and Aravind, 2010). Whereas the specific molecular and cellular functions of FAM102A family proteins are unknown, FAM102A mutations have been implicated in the progression of glaucoma (Khor et al., 2016; Li et al., 2020; Nongpiur et al., 2018; Shi et al., 2021; Zhuang et al., 2018).

Here we provide evidence in support of the model that SYM-3/FAM102A and SYM-4/WDR44 family members act together in a pathway or complex and that these proteins may carry out functions connected to intracellular trafficking. Furthermore, using a candidate-based enhancer screening approach, we have implicated ∼30 additional proteins in aECM biology in the worm embryo. We also test the model that SYM-3 and SYM-4 promote the deposition of aECM proteins during embryogenesis.

## RESULTS

### SYM-3/FAM102A and SYM-4/WDR44 colocalize within epidermal puncta

Given the previously described genetic interactions between *sym-3*, *sym-4*, and *mec-8* (Davies et al., 1999; Kelley et al., 2015; Yochem et al., 2004), we hypothesized that SYM-3 and SYM-4 may function together in a common pathway or complex or at a common subcellular location. To test this last prediction, we obtained strains expressing endogenously CRISPR-tagged SYM-3::GFP, SYM-3::mScarlet, SYM-4::GFP, and SYM-4::mScarlet and examined their expression and subcellular localization patterns. Both SYM-3 and SYM-4 were detected at low levels throughout development, with their most prominent expression observed at punctate structures along the boundary of the hyp7 epidermal syncytium and epidermal seam cells in larvae (Fig 1A, 1B, 1E, and 1F). Additional puncta were also detected within the cytoplasm of hyp7 and the seam cells, although these were less abundant. Notably, both color combinations of tagged markers revealed clear colocalization of SYM-3 and SYM-4 in the larval epidermis, consistent with these proteins acting together in cells (Fig 1 and File S1).

**Fig 1.**
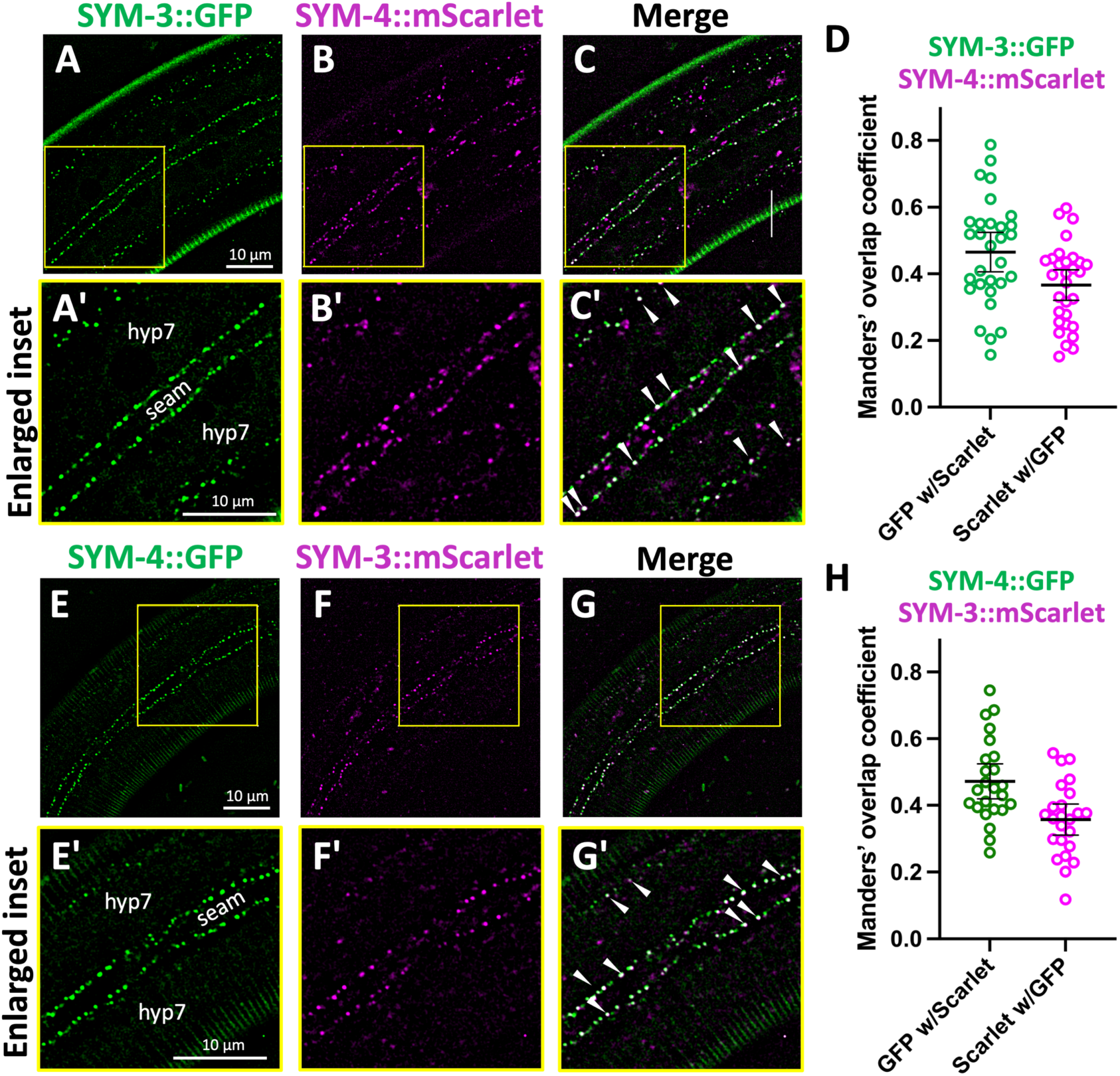
Expression and colocalization of SYM-3 and SYM-4 in the epidermis of *C. elegans* larvae.00000000. (A–C’) Localization of endogenously tagged SYM-3::GFP (A, A’) and SYM-4::mScarlet (B, B’) and a merged image (C, C’). (A’–C’) Enlarged regions from A–C as indicated by the yellow boxes. (E– G’) Localization of endogenously tagged SYM-4::GFP (E, E’) and SYM-3::mScarlet (F, F’) and a merged image (G, G’). (E’–G’) Enlarged regions from E–G as indicated by the yellow boxes. Larvae in A–G’ are at the third or fourth larval stage (i.e., L3 or L4, respectively). Locations of the seam cell and hyp7 epidermal syncytium are indicated; arrowheads indicate overlapping SYM-3–SYM-4 puncta (white dots), which are most prevalent along the seam cell–hyp7 border. (D, H) Overlap for each pair of markers from SYM-3::GFP and SYM-4::mScarlet (D) and SYM-4::GFP and SYM-3::mScarlet (H) was calculated using Manders’ overlap coefficient; mean and 95% confidence intervals (CIs) are shown in addition to individual data points. Throughout the figures, each scale bar is applicable to all similar images within a figure.

Because SYM-3 and SYM-4 play a role in anterior morphogenesis (Davies et al., 1999; Kelley et al., 2015; Yochem et al., 2004), we examined these markers in early comma-stage embryos. Although expressed at low levels, SYM-3 and SYM-4 were detected at punctate structures throughout the embryo and, like their expression in larvae, were observed to overlap in their localization (Fig S1 and File S1). As a complement to CRISPR-tagged markers, we examined the expression of *sym-3::GFP* and *sym-4::GFP* fosmid-based transgenes (Sarov et al., 2012) that were expressed from extrachromosomal arrays (Fig S1). Both transgenes rescued the Pin phenotype of corresponding *mec-8; sym-3* and *mec-8; sym-4* mutants, indicating that the protein fusions are functional. Although more variable than the CRISPR reporters, analysis of the transgenes suggested the potentially broad expression of both proteins throughout development including localization to membranes, the cytoplasm, and punctate structures (Fig S1 and File S1).

### Affinity capture proteomics implicates physical interactions between FAM102A, WDR44, and protein trafficking factors

To identify candidate protein partners of SYM-3 and SYM-4 family proteins, we carried out affinity capture proteomics using C-terminally tagged FAM102A::GFP and WDR44::GFP “bait” proteins expressed in HeLa cells (Fig 2A). Expression of both fusions was verified through western blotting and immunofluorescence, which suggested a punctate pattern in the cytoplasm of HeLa cells (Fig 2A). Two independent affinity capture proteomics experiments were carried out for each bait protein in parallel with a GFP-only control (Files S2 and S3). To narrow down the list of potential interactors, we further filtered candidates by requiring a minimum of three spectral counts (SCs) for each candidate protein as well as a ratio of bait SCs to control SCs of ≥5 (File S2). Filtered results are summarized in Fig 2B. In total we identified 333 proteins in one or both FAM102A pull-downs and 253 proteins in one or both WDR44 pull-downs.

**Fig 2.**
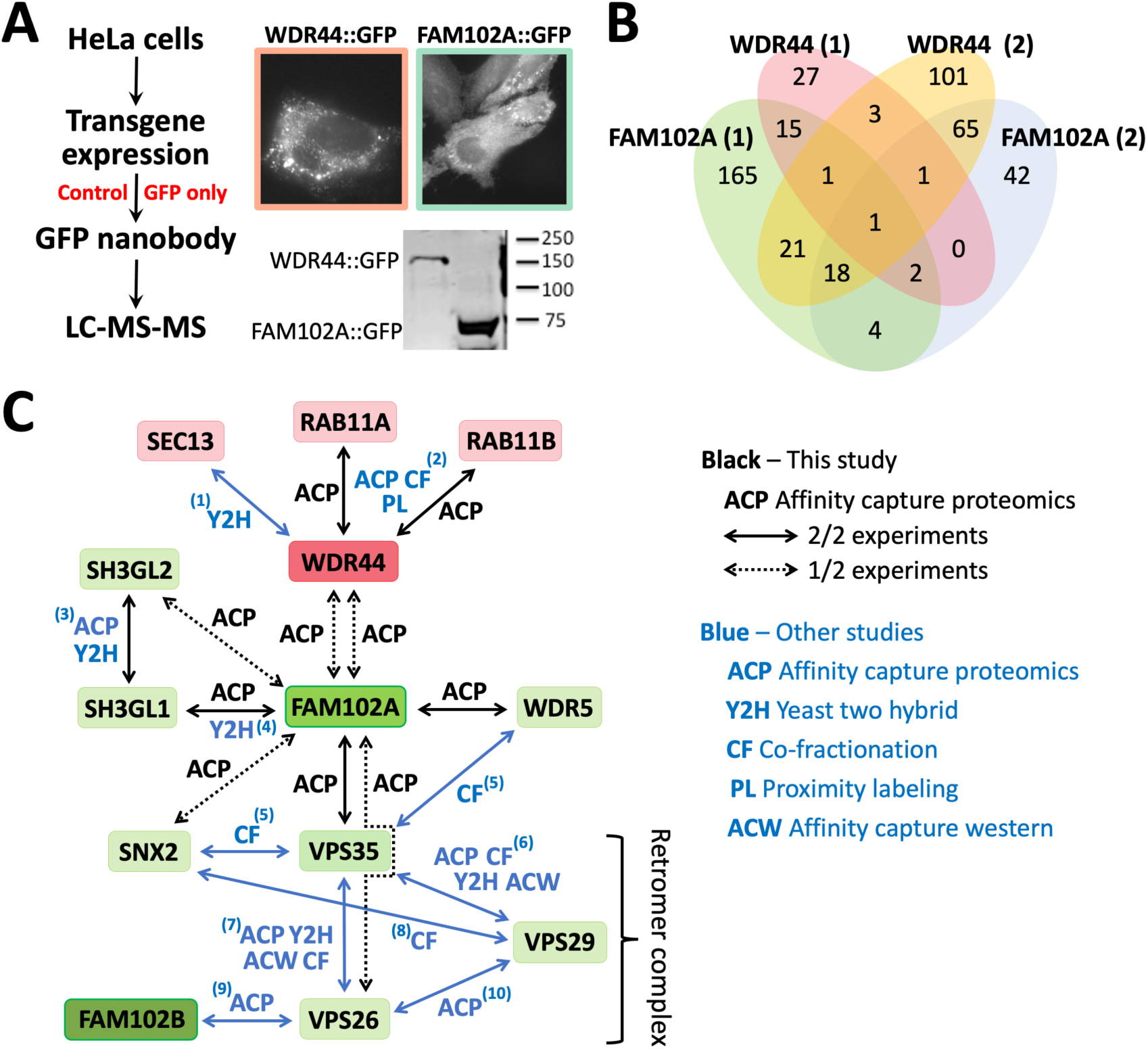
Affinity capture proteomics of the human SYM-3 and SYM-4 orthologs, FAM102A and WDR44. (A) Schematic of experimental procedure to identify FAM102A- and WDR44-interacting proteins in HeLa cells using affinity capture in conjunction with liquid chromatography and tandem mass spectrometry (LC-MS-MS) proteomics. Representative immunofluorescence (upper) and western (lower) images from HeLa cells expressing the two bait proteins fused to GFP are shown. (B) Venn diagram summarizing the results (after candidate filtering) of two independent experiments. Filtered candidates had a minimum of three spectral counts and were detected at ≥5-fold excess relative to the GFP-only control. For details see Supplemental Files S2 and S3. (C) Diagram of several highlighted interactions identified through affinity capture proteomics (black arrows, this study) as well as physical associations identified in other published studies (black arrows, this study; blue arrows, previous studies). PMID reference numbers for the published interactions are indicated by the superscript numbers. 1: 10747849 (Y2H); 2: 35271311 (ACP), 26496610 (ACP), 28514442 (ACP), 33961781 (ACP), 34079125 (ACP), 36463963 (ACP), 26186194 (PL), 10464283 (CF), 10077598 (CF); 3: 35271311 (ACP), 26186194 (ACP), 28514442 (ACP), 33961781 (ACP), 16115810 (Y2H), 25416956 (Y2H), 21900206 (Y2H); 4: 32296183 (Y2H), 25416956 (Y2H); 5: 22939629 (CF), 35831314 (CF), 22863883 (CF), 26344197 (CF); 6: 35271311 (ACP), 28514442 (ACP), 33961781 (ACP), 29893854 (ACW), 22939629 (CF), 35831314(CF), 22863883 (CF), 26344197 (CF), 32296183 (Y2H), 25416956 (Y2H); 7: 35271311 (ACP), 26496610 (ACP), 26186194 (ACP), 28514442 (ACP), 33961781 (ACP), 29893854 (ACW), 24819384 (ACW), 22939629 (CF), 35831314 (CF), 22863883 (CF), 26344197 (CF), 32296183 (Y2H); 8: 22939629 (CF), 35831314 (CF), 22863883 (CF), 26344197 (CF); 9: 26496610 (ACP); 10: 22036573 (ACP).

FAM102A was the most abundant protein identified in both FAM102A replicates (201 and 259 sequence SCs), whereas WDR44 was the most abundant protein identified in both WDR44 replicates (208 and 457 sequence SCs), indicating that the GFP pulldowns worked as expected. Moreover, FAM102A was the second-most-abundant protein identified in one of the two WDR44 replicates (19 bait SCs versus 0 control SCs), and, although enriched in the other replicate (10 bait SCs versus 5 control SCs), it failed to make the ratio cutoff for our filtered list. Correspondingly, WDR44 was identified in one of the two FAM102A replicates (5 bait SCs versus 0 control SCs). These results are consistent with both the genetic and colocalization data from *C. elegans* and suggest that FAM102A and WDR44 family proteins may function together in a conserved protein complex. We note that we were unable to validate physical interactions between SYM-3 and SYM-4 CRISPR-tagged proteins from worm lysates using immunoprecipitation–western blotting approaches, which may be due to low levels of the proteins or because our worm lysis protocols disrupted SYM-3–SYM-4 interactions.

As indicated by the Venn diagram (Fig 2B), our analysis identified proteins that were common to multiple datasets including 6 proteins present in both WDR44 datasets and 25 proteins present in both FAM102A datasets (File S2). Notably, RAB11A and RAB11B GTPases were identified in both datasets for WDR44 (Fig 2C), consistent with findings in the literature demonstrating that these proteins physically interact and function together in membrane trafficking (Mammoto et al., 1999; Thibodeau et al., 2023; Zeng et al., 1999).

Interestingly, several proteins identified in the FAM102A pulldowns are also connected to membrane trafficking including the Endophilin-A1 family members SH3GL1 and SH3GL2 (Fig 2C). Endophilins contain membrane-binding/bending BAR domains along with SH3 domains and have been implicated in several trafficking functions including clathrin-independent endocytosis (Casamento and Boucrot, 2020; Kjaerulff et al., 2011). In addition, interactions between FAM102A and two members of the retromer complex, VPS35 and VPS26, were also detected, as was an interaction with the retromer-associated protein SNX2 (Griffin et al., 2005; Gullapalli et al., 2004) (Fig 2C). Retromer is a conserved complex implicated in several endocytic processes including endosome-to-Golgi retrieval and endosome-to-plasma-membrane recycling (Seaman, 2012; Seaman, 2021; Tu and Seaman, 2021). We also note a detected interaction between FAM102A and WDR5, which is connected to the retromer complex through protein interaction data (Fig 2C). Collectively, these physical interactions suggest that WDR44 and FAM102A may perform functions connected to membrane trafficking.

### Loss of SYM-3 and SYM-4 have minimal-to-no effects on the subcellular localization of FBN-1 and NOAH-1

Given a potential role for SYM-3 and SYM-4 in the membrane trafficking of proteins, together with prior genetic interaction data (Davies et al., 1999; Kelley et al., 2015), we hypothesized that loss of either or both proteins might alter the abundance and/or localization of the matrix component FBN-1 to the aECM during embryogenesis. We obtained an endogenously tagged FBN-1::sfGFP based on a previous design (Cohen et al., 2020b) with the fluorescent marker inserted after H2408 (*fbn-1a* exon 21), just N-terminal to the extracellular ZP domain (C2436–P2671). We first detected expression of FBN-1::sfGFP in wild-type embryos just prior to the onset of morphogenesis with weak localization observed at the embryonic apical surface. Expression of aECM-associated FBN-1::sfGFP increased during early stages of embryonic morphogenesis, with expression continuing for the remainder of embryogenesis (Fig S2, Fig 3A, and Movies S1–S3).

**Fig 3.**
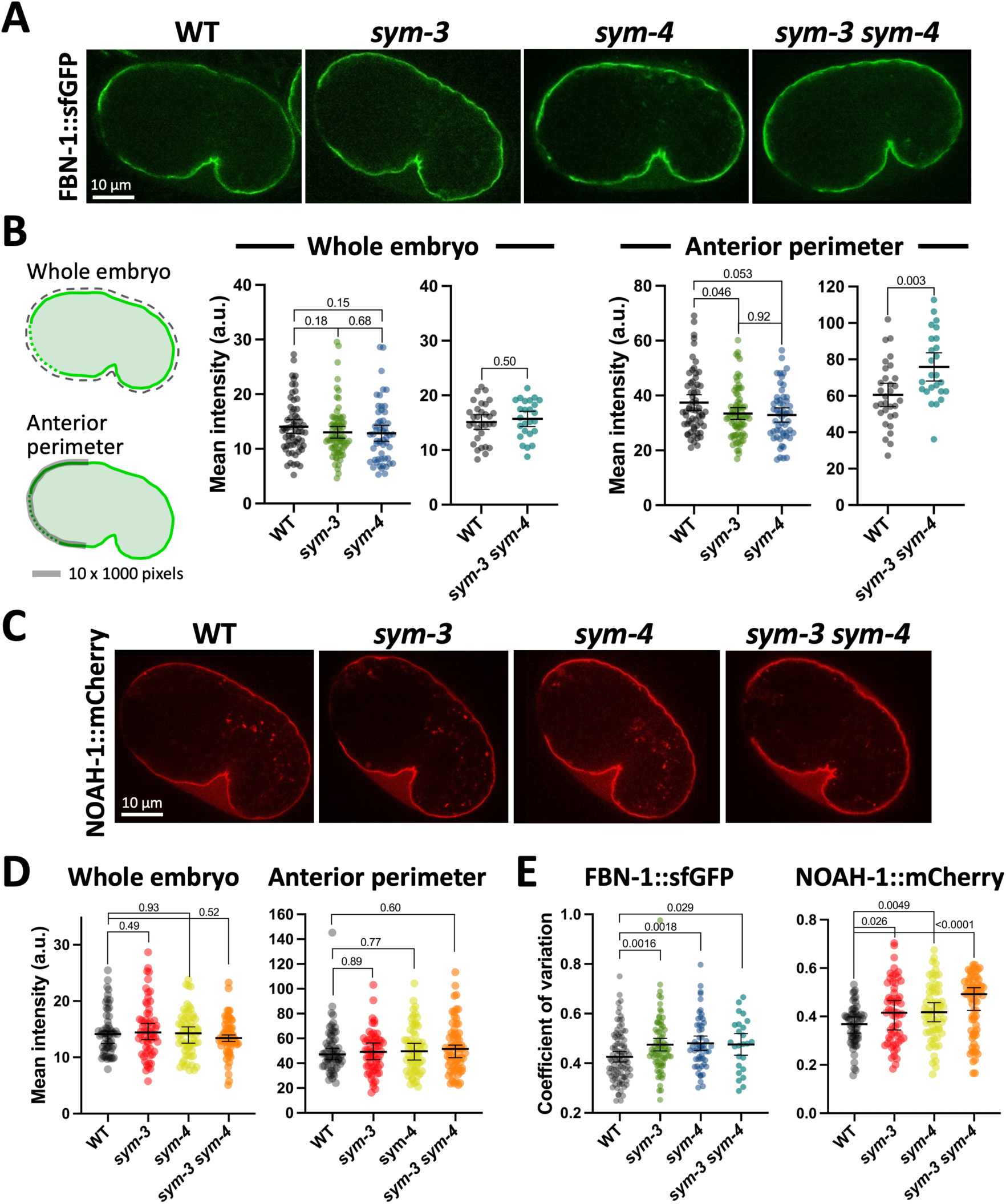
Embryonic localization of FBN-1 and NOAH-1 in *sym-3* and *sym-4* mutants. (A) Expression of FBN-1::sfGFP in early comma-stage embryos of wild type (WT) and *sym* mutants. Anterior is to the left, ventral is down. (B) Quantification of expression from genotypes shown in Panel A; leftmost diagrams of embryos indicate the regions that were quantified. (C) Expression of NOAH-1::mCherry in early comma-stage embryos of wild type and *sym* mutants. (D) Quantification of NOAH-1::mCherry as for Panel B. (E) Coefficient of variation for anterior perimeter traces. Data in B, D, and E indicate the mean ± 95% CIs in addition to individual data points. Indicated P-values (above brackets) were determined using a two-tailed Mann-Whitney test.

A preliminary visual inspection of FBN-1::sfGFP in *sym-3* and *sym-4* single mutants failed to reveal obvious differences between the mutants and wild type during morphogenesis (Fig 3A and Fig S2B). To screen for more subtle effects, we quantified expression in early comma-stage embryos, ∼370 minutes post-fertilization (at 22°C) (Chisholm and Hardin, 2005), a time point that just precedes attachment of the primordial pharynx to the epidermis (Portereiko and Mango, 2001). With respect to whole-embryo expression levels, we failed to detect a significant difference in the mean intensity of FBN-1::sfGFP in wild-type, *sym-3*, and *sym-4* embryos (Fig 3A and 3B). In addition, we examined the level of apically localized FBN-1::sfGFP in the anterior region of embryos (referred to as anterior traces), where the aECM is subsequently required to withstand pharyngeal pulling forces. Here we observed a very slight decrease (∼10–15%) in the levels of FBN-1::sfGFP in *sym* mutants, although this trend was at most marginally significant (Fig 3A and 3B). As an additional test we examined FBN-1::sfGFP in *sym-3 sym-4* double mutants. Again, no difference in mean intensities was observed for whole embryos, although we detected a slight increase (∼1.25-fold) in the anterior apical FBN-1::sfGFP signal in *sym-3 sym-4* embryos (Fig 3A and 3B). This small difference, if meaningful, runs counter to the model that SYM-3 and SYM-4 promote FBN-1 trafficking and secretion to the aECM. As a final test, we asked whether the distribution of apically localized FBN-1::sfGFP along the anterior aECM was altered in *sym* mutants by determining the coefficient of variation for FBN-1::sfGFP anterior traces. By this measure, *sym-3*, *sym-4*, and *sym-3 sym-4* mutants showed at most a modest increase (∼1.1-fold) in variability relative to wild type (Fig 3E). Taken together, these results fail to support a clear role for SYM-3 or SYM-4 in the gross deposition of FBN-1 to the aECM during embryogenesis.

We next considered the possibility that SYM-3 and SYM-4 may act on components of the aECM other than FBN-1. For this purpose, we chose to test NOAH-1, a ZP domain–containing sheath protein that acts in a mechano-transducing and structural pathway that functions in parallel to the FBN-1 (Vuong-Brender et al., 2017). Similar to the findings for FBN-1::sfGFP, however, we failed to detect substantial differences between wild-type, *sym-3*, *sym-4*, and *sym-3 sym-4* embryos based on the measures described above, although apical anterior traces of NOAH-1::mCherry suggested a slight trend toward increased variability in the mutants (∼1.1-to 1.2-fold; Fig 3C and 3D). Whereas it remains possible that SYM-3 and SYM-4 have major roles in the deposition of other aECM components, our current data do not support an obvious function for SYM-3 or SYM-4 in the deposition of NOAH-1 or FBN-1 to the aECM.

### Mis-splicing of *fbn-1* in *mec-8* mutants does not grossly alter the levels or subcellular localization of FBN-1

We previously showed that the RNA-binding protein MEC-8/RBPMS2 regulates the splicing of *fbn-1* isoforms (Kelley et al., 2015). Specifically, mutations in *mec-8* lead to the pronounced retention of sequences corresponding to introns 14–16 of *fbn-1* (Fig 4A). In testing for potential effects of *mec-8* loss on FBN-1::sfGFP expression, we were surprised to find that FBN-1::sfGFP abundance and localization to the aECM were not strongly affected in *mec-8* mutants (Fig 4B and 4C). This result was unexpected in part because intron retention often leads to the introduction of premature stop codons, leading to the degradation of mRNAs by nonsense-mediated decay and a corresponding reduction in protein levels (Longman et al., 2008; Lykke-Andersen and Jensen, 2015). If anything, however, we observed a slight increase (∼1.25-fold) in the total abundance of FBN-1::sfGFP as well as a very slight increase (∼1.15-fold) in the level of FBN-1::sfGFP along the anterior perimeter of early comma-stage embryos (Fig 4B and 4C). Given that the Pin phenotype of *mec-8; sym-4* mutants can be largely rescued by the expression of a single *fbn-1* cDNA isoform (*fbn-1e*) (Kelley et al., 2015), this suggests that the underlying defect of FBN-1 in *mec-8* mutants is due to altered functional properties of the protein rather than a reduction in total levels of the protein or defects in its targeting to the aECM.

**Fig 4.**
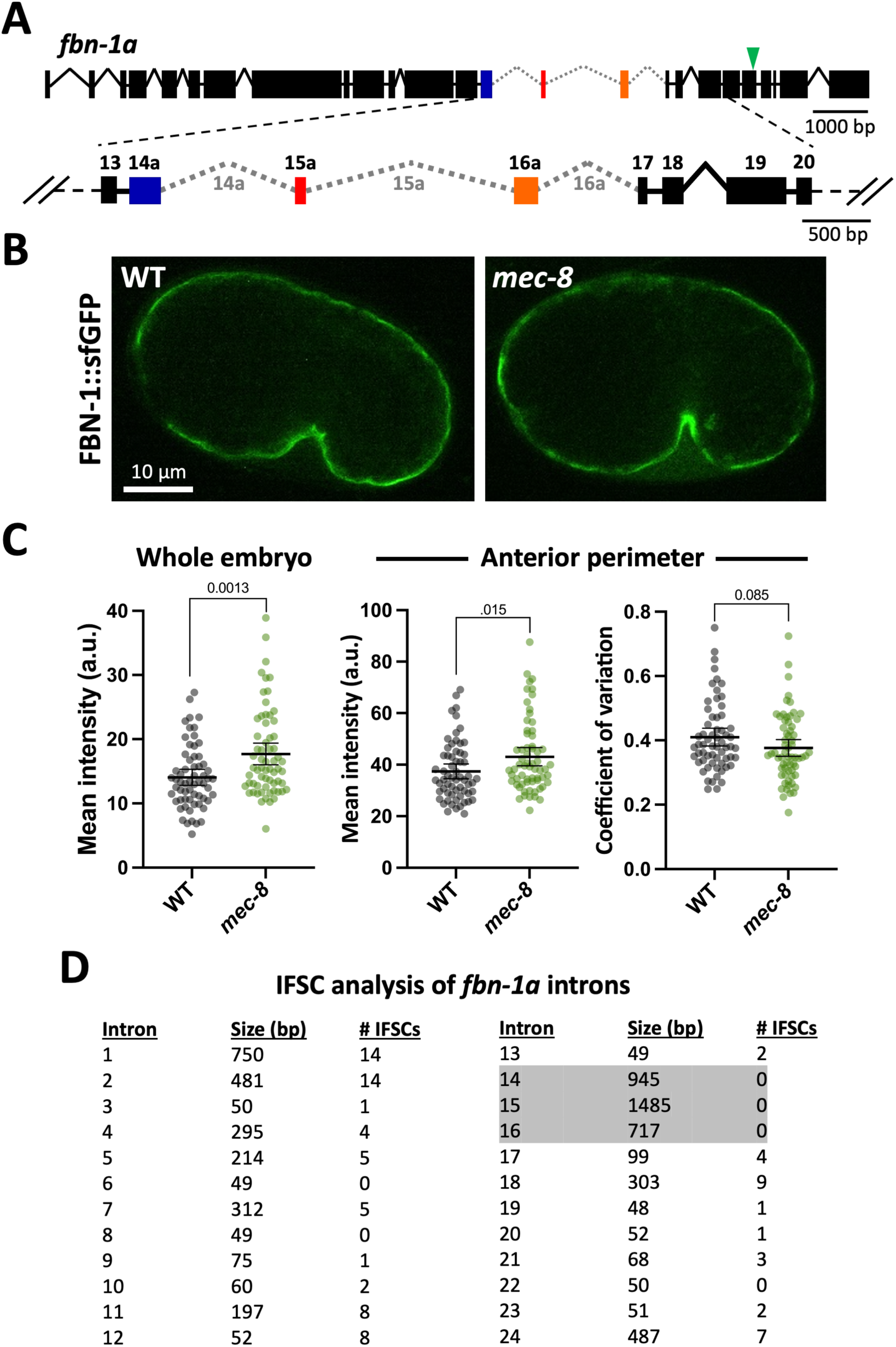
Loss of MEC-8 does not strongly affect the embryonic expression or localization of FBN-1. (A) Diagram of the *fbn-1* genomic locus (isoform ‘a’) with exons and introns regulated by MEC-8 (exons 14–16) highlighted in color and indicated by gray dotted lines, respectively. Green arrowhead in upper diagram indicates the sfGFP insertion site. (B and C) Expression (B) and quantification (C) of FBN-1::sfGFP in early comma-stage embryos of wild type and *mec-8* mutants (as described for Fig 3A, 3B, and 3E). Data are shown as the mean ± 95% CIs in addition to individual data points. Indicated P-values (above brackets) were determined using a two-tailed Mann-Whitney test. (D) The size and the number of in-frame stop codons (IFSCs) are indicated for the 24 introns of *fbn-1a*. Note the striking absence of IFSCs in introns 14–16 (shaded).

The above result may be partially explained by the absence of any in-frame stop codons in sequences corresponding to *fbn-1* introns 14–16 (Fig 4D; WormBase) (Stein et al., 2001). Moreover, RNA sequencing (RNAseq) data available on WormBase indicate that introns 14–16, although present at relatively low levels in wild-type worms, are retained at higher frequencies than *fbn-1* intronic sequences that are not regulated by MEC-8 (Fig S3A; WormBase). Together, these findings suggest that introns 14–16 may be more accurately categorized as alternate exons, which are retained to some extent even within wild-type (MEC-8 competent) worms. Consistent with this, *fbn-1* was not identified as a target of endogenous nonsense mediated decay in *C. elegans* (Muir et al., 2018), suggesting that these elongated *fbn-1* species may serve some physiological function.

As a follow up, we were interested to know if sequences corresponding to *fbn-1* introns 14–16 are conserved. In examining the peptide sequences encoded by these regions, we failed to identify any known motifs, although one region within intron 16 is predicted to encode a 190-aa peptide segment associated with amidase domain–containing proteins (7.59e-05; based on an NCBI–CD Search). BLAST searches for related peptide sequences in other *Caenorhabditis* species failed to detect significant similarities. Furthermore, predicted gene structures of *fbn-1* orthologs in other *Caenorhabditis* species did not indicate strong conservation of exon–intron boundaries (WormBase). Nevertheless, we did find one example of a potential alternative exon corresponding to intron 14 of *Caenorhabditis briggsae fbn-1*. This region consists of a 1095-bp predicted intronic sequence that, based on the available data, contains no in-frame stop codons and forms a continuous open reading frame with the adjacent upstream and downstream exons (Fig S3B and S3C). These observations suggest that alternative exons in *fbn-1* homologs— and possibly other genes—may have escaped previous detection due to gene prediction models and/or low representation of these sequences within wild-type mRNA pools.

### A focused RNAi-enhancer screen for the Pin phenotype implicates new factors involved in anterior morphogenesis and force resistance

Recent studies have implicated multiple proteins that contribute to the functions and mechanical properties of the aECM during development (Cohen et al., 2020a; Cohen et al., 2019; Cohen et al., 2020b; Cohen and Sundaram, 2020; Forman-Rubinsky et al., 2017; Heiman and Shaham, 2009; Kelley et al., 2015; Li Zheng et al., 2020; Vuong-Brender et al., 2017). These include FBN-1, DEX-1, and DYF-7, all three of which function in the resistance of the anterior epidermis to pharyngeal pulling forces (Cohen et al., 2019; Kelley et al., 2015). To identify novel aECM components and regulators, we carried out a directed RNAi screen focusing on gene categories often associated with aECM proteins (Cohen and Sundaram, 2020). In total we screened 425 genes whose products were indicated to be secreted (Suh and Hutter, 2012) or glycosylated (Fan et al., 2005; Kaji et al., 2007; Olsen and Mann, 2013) or that oscillate during larval molting cycles (Hendriks et al., 2014; Meeuse et al., 2020), with most tested genes matching at least two of these criteria (File S4). Specifically, we tested for enhancement of the larval-lethal Pin phenotype in *lin-35; sym-4* mutants by RNAi-feeding methods, in which the *lin-35* mutation served to increase the sensitivity to somatic RNAi (Lehner et al., 2006; Wang et al., 2005). For reference, whereas 8–11% of *lin-35; sym-4* mutants grown in the presence of the control RNAi vector displayed the Pin phenotype, *fbn-1(RNAi)* led to expression of the Pin phenotype in ∼97% of larvae (Fig 5 and File S4), consistent with our previous findings (Kelley et al., 2015).

**Fig 5.**
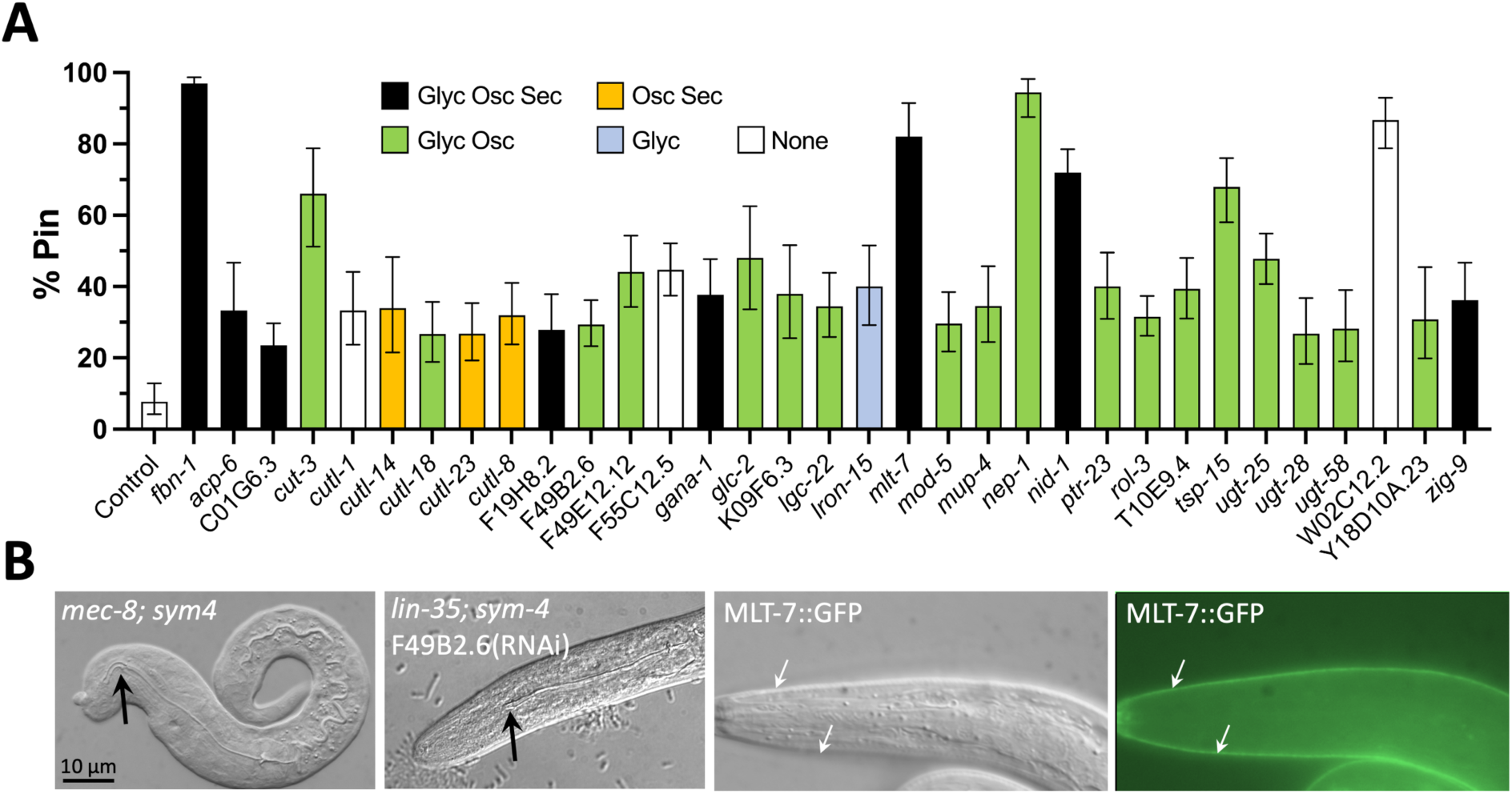
RNAi enhancers of the Pin phenotype. (A) Percentage of F1 progeny displaying the Pin (pharynx-ingressed) phenotype after growth of *lin-35(n745); sym-4(mn619)* P0 worms on the indicated RNAi feeding clones. Bars are color coded according to whether their predicted gene products are glycosylated (Gly) and/or oscillating (Osc) and/or secreted (Sec); white bars indicate that none of these categories were met. File S4 contains raw data for two experimental replicates along with category assignments for all genes tested. Results for replicate #2 are shown in Panel A. Error bars represent the 95% CIs. Based on a (pairwise) Fisher’s exact test relative to the vector control, all displayed clones had a P-value of <0.0001. (B) Two leftmost panels show representative images of a *mec-8 (u74); sym-4 (mn619)* and a *lin-35; sym-4; F49B2.6(RNAi)* L1 larva displaying the Pin phenotype. Black arrows indicate the posteriorly displaced terminus of the internalized/ingressed pharynx. Two rightmost panels show a fluorescence and corresponding DIC image of an L1 larva expressing full-length MLT-7::GFP from an extrachromosomal array (*fdEx310*), which localizes to the aECM (white arrows).

Of the 425 genes tested, we identified 32 that reproducibly increased the Pin phenotype of *lin-35; sym-4* mutants to levels that were at least 2-fold higher than that of the vector control (Fig 5A and File S4). Not surprisingly, we identified a number of genes encoding proteins with known or proposed functions in aECM biology including sheath and/or cuticle structural components such as NID-1/nidogen 1 (Kang and Kramer, 2000); the ZP domain–containing protein CUT-3 (Sapio et al., 2005); and the cuticulin-like proteins CUTL-1, -8, -14, -18, and -23, about which little is known. In addition, the screen identified candidate aECM modifiers including the peroxidase collagen cross-linker MLT-7 (Thein et al., 2009). Consistent with a role for MLT-7 in the aECM, a fosmid-based MLT-7::GFP reporter localized to the larval cuticle (Fig 5B). Other candidate aECM regulators included the receptor tyrosine kinase ROL-3 (Jones et al., 2013); the neprilysin-family peptidase NEP-1 (Spanier et al., 2005), an ortholog of the human endoplasmic reticulum peptidase F49B2.6/ERAP1; and a Patched-related protein, PTR-23, implicated in molting and membrane integrity (Choi et al., 2016; Zugasti et al., 2005). In addition, the screen identified several uncharacterized proteins as well as proteins with conserved motifs but with no previous connection to aECM biology.

We note that of the 32 RNAi clones that tested positive in our assay for Pin enhancement, only two had been previously shown to affect larval development in RNAi-feeding screens using RNAi-hypersensitive strains harboring either the *lin-35* or *rrf-3* mutations (File S4) (Ceron et al., 2007; Simmer et al., 2003). As such, the *lin-35; sym-4* background may be useful for identifying and testing additional candidate aECM regulators and components.

### Loss of *mir-51* microRNA family members causes a Pin phenotype that shows differences from *mec-8; sym* double mutants

As an alternative means of identifying genes involved in anterior morphogenesis and force resistance, we examined the literature for evidence of previously reported Pin-like phenotypes. Notably, deletion of all six members of the *mir-51* family of microRNAs (miRNAs; *mir-51–56*) leads to abnormal pharyngeal ingression, although this phenotype differs from that of other aECM mutants in its morphology and because of the presence of detached cell-like bodies (referred to hereafter as “cell bodies”) proximal to the nascent buccal cavity (Fig 6A–C) (Shaw et al., 2010).

**Fig 6.**
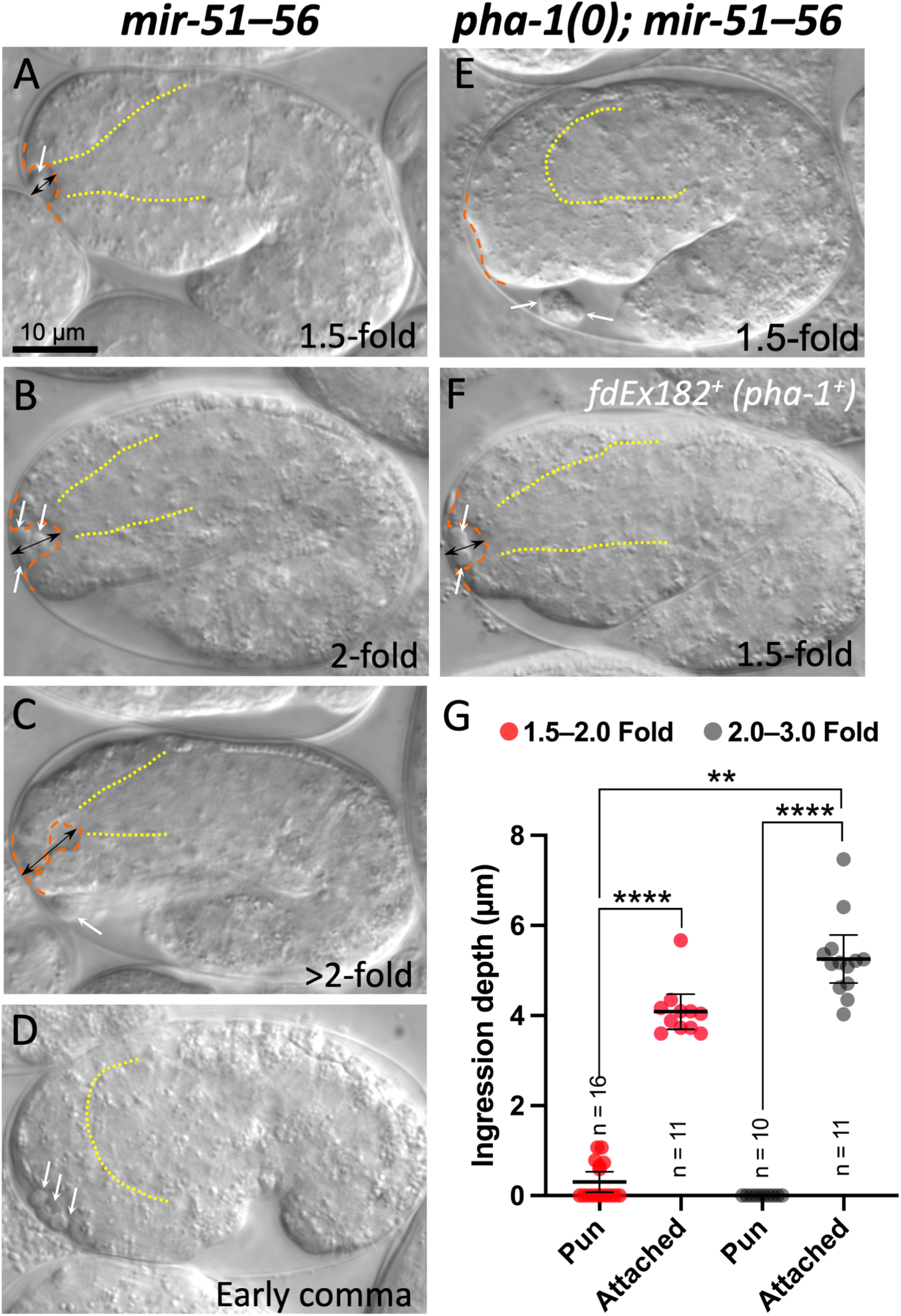
Loss of *mir-51* family members leads to a Pin phenotype that is distinct from that of *mec-8; sym* mutants. (A–F) Representative *mir-51–56* (A–D) and *pha-1(0); mir-51–56* (E, F) embryos. Anterior is to the left, ventral is down. Dashed orange lines trace the anterior-most regions of embryos including the nascent buccal cavity; dotted yellow lines indicate the basement membrane surrounding the primordial pharynx. In *mir-51–56* embryos in which the pharynx has attached to the nascent buccal cavity (A–C), an abnormal anterior ingression (doubled-headed black arrows; Pin phenotype) is observed in embryos at the ∼1.5-to 2.0-fold stages. In *mir-51–56* early comma-stage embryos (D), prior to pharyngeal attachment, no pharyngeal ingression is observed. Likewise, in 1.5-fold-stage embryos in which pharyngeal attachment has been prevented by loss of *pha-1* (E), ingression is also not observed, demonstrating that pharyngeal attachment is required for anterior ingression in *mir-51–56* mutants. Also note detached cell bodies (white arrows) in all embryos including those in which the pharynx has not attached. Panel F depicts a *pha-1(0); mir-51–56* embryo that expresses wild-type *pha-1* from a transgenic array (*fd182^+^*), leading to pharyngeal attachment and abnormal anterior ingression. (G) Ingression depths were measured as indicated by the double-headed arrows (shown in A–C, and F). Depths were measured in *pha-1; mir-51–56* mutants in which *pha-1* was rescued (*fdEx182*) versus those in which *pha-1* was not rescued (*fdEx182^−^*). Mean and 95% CIs are shown as well as individual data points. P-values are based on a two-tailed Mann-Whitney test. ****P < 0.0001, **P < 0.01.

To determine if the Pin phenotype of *mir-51–56* mutants is due to an inability to withstand pharyngeal pulling forces, we generated *mir-51–56* mutant strains in which pharyngeal attachment was prevented by loss of *pha-1*. Approximately 85% of *pha-1* null mutants fail to establish a connection between the pharynx and adjacent anterior cells of the forming buccal cavity (Fay et al., 2004; Kuzmanov et al., 2014), and loss of this connection strongly suppresses pharyngeal ingression defects in *mec-8; sym-3* double mutants (Kelley et al., 2015). Notably, whereas 21 of 22 *mir-51–56* mutant embryos exhibited the Pin phenotype, only 5 of 38 *pha-1; mir-51–56* mutants in which the pharynx failed to attach (the Pun phenotype) showed anterior ingression defects (Fig 6E). Moreover, restoring a pharyngeal connection in *pha-1; mir-51–56* mutants by expression of a wild-type *pha-1^+^* transgene (*fdEx182*) (Granato et al., 1994) led to ingression defects in 25 of 25 embryos (Fig 6F).

To further quantify these effects, we measured ingression depths in *pha-1; mir-51–56* embryos in which pharyngeal attachment occurred (*fdEx182^+^– pha-1^+^*) versus those in which it had failed (*fdEx182^−^– pha-1^−^*). Whereas 1.5-to 2-fold-stage embryos in which attachment failed had an average ingression depth of 0.31 µm, sibling pharynx-attached embryos had an average ingression depth of 4.1 µm, which increased to 5.3 µm at the 2-to 3-fold stage (Fig 6G). Our data are consistent with pharyngeal pulling forces being an underlying cause of the Pin phenotype in *mir-51–56* mutants. Moreover, the finding that *mir-51* family members regulate cadherins (Shaw et al., 2010) suggests that cell–cell adhesion may contribute to the resistance of pharyngeal pulling forces by anterior epithelial cells.

We were also interested to determine if pharyngeal pulling might contribute to the cell-body detachment phenotype of *mir-51–56* mutants. In this case we failed to observe any difference between the percentage of embryos containing detached cell bodies in *pha-1; mir-51–56* embryos (98%; n = 50) versus *pha-1; mir-51–56; fdEx182* embryos that express the *pha-1^+^* transgene (97%; n = 79). In addition, cell detachment could be observed in some early comma-stage *mir-51–56* embryos, a time point that precedes pharyngeal attachment (Fig 6D). Thus, whereas the ingression phenotype of *mir-51–56* mutants can be attributed to pharyngeal pulling forces, the cell detachment phenotype appears to occur independently of pharyngeal pulling. Collectively, these data implicate miRNAs and their targets in the regulation of anterior morphogenesis and the resistance to pharyngeal pulling forces.

## Discussion

Genetic, in vivo expression, and proteomic data are consistent with the model that SYM-3/FAM102A and SYM-4/WDR44 function together in a shared pathway or complex. This includes previous genetic interaction data in *C. elegans* (Davies et al., 1999; Kelley et al., 2015; Yochem et al., 2004) along with new data showing that SYM-3 and SYM-4 colocalize to cytoplasmic and membrane-associated puncta (Fig 1 and Fig S1). Although the identity of these puncta are unknown, they are reminiscent of vesicles or compartments associated with membrane trafficking. In addition, our proteomic data in mammalian cells (Fig 2, Files S2 and S3) are consistent with WDR44 and FAM102A family proteins functioning together in a complex and potentially acting with additional trafficking components such as RAB11, endophilins, and the retromer complex.

WDR44 was first identified as a binding partner and putative effector of RAB11 (Mammoto et al., 1999; Zeng et al., 1999). These studies implicated WDR44 in transferrin recycling and cell migration, consistent with the functions of RAB11 (Bhuin and Roy, 2015; Chung et al., 2014; Gibieza and Petrikaite, 2021; Ramel et al., 2013; Welz et al., 2014). WDR44 associates specifically with the activated GTP-bound form of RAB11 and is colocalized with RAB11 at perinuclear regions (Mammoto et al., 1999; Zeng et al., 1999). Subsequent work has shown that WDR44 and RAB11 also function together in the control of cilium growth and that their interaction is controlled by Akt phosphorylation of WDR44 (Goldenring, 2019; Walia et al., 2019). Although these studies may suggest a potential role for SYM-4 in conjunction with *C. elegans* RAB-11 proteins, a recent study characterizing the peptide domains responsible for WDR44–RAB11 binding identified a key region within WDR44 that is not conserved in SYM-4 (Thibodeau et al., 2023).

WDR44 also binds Sec13 (Mammoto et al., 2000), a component of COPII vesicles required for budding from the endoplasmic reticulum (Gillon et al., 2012; Lord et al., 2013; Roberg et al., 1997). More recently, WDR44 was found to promote the export of newly synthesized E-cadherin to the plasma membrane in mammalian cells through its interactions with GRAF1b/2 (Lucken-Ardjomande Hasler et al., 2020), raising the possibility that SYM-4 could contribute to anterior morphogenesis through the regulation of cadherins. Notably, loss of *mir-51* family members leads to cadherin misregulation along with associated defects in anterior morphogenesis (Shaw et al., 2010), including the Pin phenotype (Fig 6).

Relatively little is known about the molecular functions of mammalian FAM102A/EE1G1, although protein interaction data from our work and other high-throughput studies suggest possible roles in trafficking (Fig 2, Files S2 and S3). SYM-3/FAM102A contains an NT-C2 membrane localization domain that is found in several trafficking proteins (Shi et al., 2010; Wang et al., 2016; Zhang and Aravind, 2010). Moreover, a study in worms and flies implicated *sym-3* and its *Drosophila melanogaster* ortholog, CG8671, along with several endocytic regulators, in the uptake of dsRNA molecules during RNA silencing (Saleh et al., 2006). Finally, recent reports have implicated FAM102A variants in glaucoma (Khor et al., 2016; Li et al., 2020; Nongpiur et al., 2018; Shi et al., 2021; Zhuang et al., 2018), an eye disorder linked to altered ECM components and abnormal ECM histology (Hernandez and Ye, 1993; Nikhalashree et al., 2019; O’Callaghan et al., 2017; Tribble et al., 2018; Vranka et al., 2015; Wiggs, 2015). Thus, FAM102A family members may carry out conserved functions in trafficking and ECM biology.

Despite the above inferences, our results do not support the straightforward model that SYM-3 and SYM-4 promote the trafficking or deposition of sheath-associated aECM proteins during embryogenesis (Fig 3). As such, further studies in *C. elegans* and other systems will be necessary to obtain a clearer picture of the cellular and developmental functions of both FAM102A and WDR44 family proteins. Nevertheless, our studies were successful in identifying new regulators of anterior morphogenesis (Fig 5 and File S4), many of which likely affect the structure and functions of the embryonic aECM. In addition, we obtained somewhat unexpected results regarding the role of the conserved MEC-8/RBPMS splicing factor in the regulation of FBN-1; loss of MEC-8 likely leads to the increased incorporation of previously unannotated alternative exons (Fig 4) (Kelley et al., 2015). Collectively, our findings help to lay the groundwork for future investigations into roles and regulation of the apical ECM during development.

## MATERIALS AND METHODS

### Fluorescent reporters

Strains expressing endogenously tagged reporters include PHX1529 [*sym-3::GFP(syb1529)*], PHX4755 [*sym-3::mScarlet(syb4755)*], PHX1532 [*sym-4::GFP(syb1532)*], PHX4831 [*sym-4::mScarlet(syb4831)*], and PHX1726 [*fbn-1::sfGFP(syb1726)*], which were designed by DSF and generated by SUNY Biotech with fluorescence tag insertions in *sym-3* and *sym-4* occurring before the stop codon; *sfGFP* was inserted after H2408 (*fbn-1a* exon 21). Colocalization studies (Fig 1 and Fig S1) were carried out using WY1972 [*sym-3::GFP(syb1529); sym-4::mScarlet(syb4831)*] and WY1973 [*sym-4::GFP(syb1532); sym-3::mScarlet(syb4755)*]. NOAH-1::mCherry (*mc68*) was generated as described (Vuong-Brender et al., 2017). SYM-3::GFP (WY1012; *fdEx244*) and SYM-4::GFP (WY1026; *fdEx241*) extrachromosomal array lines were generated by injecting recombineered fosmids containing the *sym-3* and *sym-4* genomic loci with GFP fused to the C termini (Sarov et al., 2012), together with pRF6 (*rol-6gf*) (Mello et al., 1991; Sarov et al., 2012). MLT-7::GFP (WY1313; *fdEx310*) was generated by injecting a recombineered fosmid containing the *mlt-7* genomic locus fused to GFP at the C terminus (Sarov et al., 2012) together with rescuing sequences for *unc-119* into *unc-119(ed3)* (Maduro and Pilgrim, 1996) mutants to generate the extrachromosomal array *fdEx310*.

### Imaging

Confocal fluorescence images in Figs 1A–C, 1E–G, 3A, 3C, 4B, S1A–C and S1F, and S2 and Movies S1–S3 were obtained using an Olympus IX83 inverted microscope coupled with an Olympus SPINSR W1 SoRa Spinning Disk Confocal microscope. A 100×, 1.35 N.A. silicone oil objective was used to obtain the z-stack images. cellSens Dimension v3.1 built-in software (Olympus) was used for image acquisition with appropriate filters (GFP and RFP). Images in Fig 5B and 6A–F were obtained using a Cannon compound microscope (100×) with DIC and fluorescence capabilities. Images were pseudocolored for presentation.

### Colocalization analysis

A Wiener deconvolution algorithm in the cellSens Dimension v3.1 software was used to deconvolute the raw z-stack images. The required z-plane was extracted from both raw z-stack and deconvoluted z-stack images. The deconvoluted images were then filtered using the Gaussian filter option (radius = 10 pixels), and the resulting filtered images were then subtracted using the image calculator function from the original deconvoluted image. Subsequently, filter-subtracted images were subjected to a thresholding function to obtain thresholded binary images to be used as masks. The thresholded binary masks were then combined with the background-subtracted raw images (rolling ball radius algorithm; radius = 50 pixels) using the “AND” Boolean operation. Finally, the polygon tool was used to identify the region of interest (e.g., seam cell region) in the combined images. The Coloc2 plugin was used to calculate the colocalization between both green and red channels based on the Manders’ coefficient. Apart from the deconvolution, the entire colocalization analysis was performed using NIH Fiji software.

### Expression and purification of anti-GFP nanobody

The anti-GFP nanobody was purified as described (Topalidou et al., 2016). Briefly, the bacterial expression vector pLaG16 was transformed in Arctic Express (DE3) cells (Agilent). Bacteria were then induced with 0.1 mM IPTG for 16 hours at 8°C and centrifuged at 5,000 *g* for 10 minutes at 4°C. Cells were incubated for 1 hour in TES buffer (0.2 M Tris-HCl, pH 8; 0.2 mM EDTA; 0.5 M sucrose) and spun down for 10 minutes at 6,000*g* at 4°C, and the pellet was resuspended in TE buffer (0.2 M Tris-HCl, pH 8; 0.5 mM EDTA). The resuspended cells were then incubated at 4°C for 45 minutes and centrifuged at 30,000*g* at 4°C for 30 minutes. The supernatant (periplasmic fraction) was bound to PerfectPro Ni-NTA Agarose affinity resin (5 Prime Sciences) for 1 hour at 4°C. The resin was then washed once with wash buffer I (20 mM sodium phosphate, pH 8.0; 0.9 M NaCl) and twice with wash buffer II (20 mM sodium phosphate, pH 8.0; 150 mM NaCl; 10 mM imidazole). Elution was carried out using His elution buffer (20 mM sodium phosphate, pH 8.0; 150 mM NaCl; 250 mM imidazole), and the eluent was dialyzed with PBS. Recombinant anti-GFP nanobody was conjugated to epoxy-activated magnetic Dynabeads M270 (Life Technologies) using 10 μg recombinant protein/1 mg Dynabeads. Conjugations were carried out for 18 h in 0.1 M sodium phosphate, pH 8.0 and 1 M ammonium sulfate at 30°C. Beads were washed once with 100 mM glycine, pH 2.5; once with 10 mM Tris, pH 8.8; four times with PBS; and twice with PBS plus 0.5% Triton X-100. Beads were stored at –20°C in PBS with 50% glycerol.

### Cell culture

HeLa cells were grown at 5% CO2 at 37°C in Dulbecco’s Modified Eagle Medium (DMEM) supplemented with 10 % fetal bovine serum (FBS) and 1% penicillin/streptomycin (P/S). Cells were passaged every 2–3 days when confluent. Cells were transfected when needed with 15 μg plasmid using Lipofectamine 2000 (ThermoFisher).

### Immunostaining of HeLa cells

2x10^5^ HeLa cells per well were seeded onto UV sterilized cover slips placed in 12-well cell culture plates. 24-48 hours later, cells were rinsed three times with ice-cold PBS and fixed with 4% paraformaldehyde in PBS for 20 minutes at room temperature. After fixation, cells were rinsed twice with PBS, permeabilized with 0.5% Triton X-100 in PBS for 5 minutes and washed again twice with PBS. Cells were then placed in 5% milk in PBS for 1 hour, followed by staining with mouse monoclonal anti-GFP (1:300, Santa Cruz #sc-9996) in 0.5% milk in PBS for 1 hour at room temperature. The cells were then washed with PBS three times (5 minutes each) and incubated with Alexa Fluor 488 anti-mouse secondary antibody (1:1000, Jackson Immunoresearch #115-545-146) at 4°C overnight. The cells were washed with PBS three times for 5 min each and examined by fluorescence microscopy (Nikon 80i wide-field compound microscope).

### Protein extraction and immunoblotting

For protein extraction and immunoblotting cells were lysed as described below (Mass spectrometry). 40 µl of the lysed supernatant was resuspended in 6x Laemmli loading buffer. Samples were resolved on 10% SDS-polyacrylamide gels and blotted onto nitrocellulose membranes. Membranes were blocked using 3% milk in TBST (50 mM Tris, pH 7.4, 150 mM NaCl, 0.1% Tween 20) for 1 h at room temperature and stained with mouse monoclonal anti-GFP (1:2000, Santa Cruz #sc-9996) in 3% milk in TBST at 4°C overnight, followed by three 5-min washes in TBST. Membranes were then stained with Alexa Fluor 680–conjugated goat anti-mouse antibody (1:20,000; Jackson Laboratory; #115-625-166) in 3% milk in TBST at room temperature, followed by three 5-min washes in TBST. Images were developed using a LI-COR processor.

### Expression constructs

The WDR44::GFP construct was constructed by amplifying human WDR44 cDNA from a HeLa cDNA library using primers: 5’–gactcagatctcgagctcaagcttcgatggcgtcggaaagc–3’ and 5’– ctcaccatggtggcgaccggtggatcagatacattttttcttttattaacaaacac–3’. The FAM102A::GFP construct was constructed by amplifying human FAM102A cDNA from a HeLa cDNA library using primers: 5’– gactcagatctcgagctcaagcttcgatggctttcttgatgaagaaga–3’ and 5’–ctcaccatggtggcgaccggtggatcatggctttcaatcacaactgg–3’. The PCR products were inserted in pEGFP-N1 vectors digested with EcoRI/BamHI using Gibson cloning.

### Mass spectrometry sample preparation

For immunoprecipitation, 4 × 10^6^ HeLa cells were plated onto 10-cm Petri dishes. Twenty-four hours later, cells were transfected with either FAM102A::GFP, WDR44::GFP, or GFP. After 24 hours, cells were washed with PBS and harvested in 400 μL lysis buffer (50 mM Tris, pH 7.5; 150 mM NaCl; 1% NP-40; Protease inhibitor cocktail [Pierce]). Lysates were passed 10 times through a 20-G needle and incubated for 30 minutes at 4°C. Lysates were then pre-cleared by centrifugation at 20,000*g* for 15 minutes at 4°C, and the supernatant was incubated with 30 μg anti-GFP nanobody bound to magnetic beads for 2 hours at 4°C. The beads were washed three times with 250 μL wash buffer (50 mM Tris, pH 7.5; 300 mM NaCl; 1% NP-40). Samples were eluted in two steps with (1) 0.2 M glycine (pH 2.5) and (2) 1 M NH_4_OH. Eluents were neutralized with 0.33 M Tris-HCl (pH 7.5) and stored at –80°C.

### Mass Spectrometry

Mass spectrometry was carried out at the University of Washington Proteomics Resource (Seattle, WA). Peptide digests were analyzed with an Orbitrap Fusion Tribrid Mass Spectrometer (Thermo Fisher Scientific). Peptides were separated online by reversed-phase chromatography on a nanoACQUITY UPLC system (Waters, Milford, MA) using a heated C18 column over a 120-minute gradient. The mass spectrometer was operated in data-dependent acquisition mode. Full survey MS scans were acquired in the Orbitrap at a resolution of 120,000 at 400 m/z with a scan range of 375 to 1575 m/z. MS/MS scans were acquired on selected precursor ions which were fragmented using higher-energy collisional dissociation fragmentation with a 29% collision energy and a stepped collision energy of 5%, detected at a mass resolution of 15,000 at 400 m/z in the Orbitrap, with a maximum injection time of 60 ms and an automated gain control target of 30,000. The MS/MS spectra were searched against the UniProt human protein sequence database using the Comet search engine with the following settings: fully tryptic allowing up to two missed cleavages, 20 ppm precursor tolerance, oxidized methionine as a variable modification, and carbamidomethylated cysteine as a static modification. The peptide-spectrum matches were analyzed by the PeptideProphet tool and those results uploaded to the Mass Spectrometry Data Analysis Platform (MSDaPl) for protein inference and subsequent analysis.

### Data files

Data resulting from the proteomic studies, including filtered and peptide-read data, are contained in Files S2 and S3, respectively. Data presented in File S2 were analyzed using the Mass Spectrometry Data Platform (MSDaPl) (Sharma et al., 2012). Identified candidates were filtered by requiring a minimum of three spectral counts (SCs) for each candidate protein as well as a ratio of bait SCs to control SCs of ≥5.

### Mutant strains

Analyses of FBN-1::sfGFP and NOAH-1::mCherry in *sym* mutants (Fig 3) were carried out using strains WY2047 [*sym-3(tm6634); mc68[noah-1::mCherry*], WY2048 [*sym-4(tm6616); mc68[noah-1::mCherry*], WY2063 [*sym-3(tm6634) sym-4(tm6616); mc68[noah-1::mCherry*], WY1978 [*sym-4(tm6616); fbn-1::sfGFP (syb1726)*], WY1979 [*sym-3(tm6634); fbn-1::sfGFP (syb1726)*], and WY2062 [*sym-3(tm6634) sym-4(tm6616); fbn-1::sfGFP (syb1726)*]. Analyses of FBN-1::sfGFP in *mec-8* mutants (Fig 4) was carried out using WY2028 [*mec-8(e398); fbn-1::sfGFP (syb1726)*]. The Pin enhancer screen (Fig 5) was carried out using WY964 [*lin-35(n745); sym-4(mn619)*]. The *mir-51–56* analysis (Fig 6) was carried using SX356 [*nDf67, mir-52(n4114); nDf58; mjEx123 (mir-52^+^ rescuing transgene + DLG-1∷mCherry)*] (Shaw et al., 2010) and WY884 [*pha-1(tm3671); nDf67, mir-52(n4114); nDf58; mjEx123; fdEx182 (pha-1^+^ rescuing transgene + SUR-5::GFP)*].

### FBN-1 and NOAH-1 reporter analyses

Both anterior perimeter and whole worm mean intensity analyses were performed using Fiji software. Desired planes were selected from the z-stack of raw embryo images. A segmented line tool was used to trace the anterior perimeter of each embryo; the width of the traced line was 10 pixels. The length of the trace was set to be the same for all embryos (1000 pixels). Mean intensity data were acquired from anterior perimeter traces and plotted. Meanwhile, a coefficient of variation (CV) analysis was performed using the anterior perimeter mean intensity data with the following formula: CV = SD/mean (i.e., the SD of each embryo was divided by its corresponding mean). Movies were created from the z-stack images using Fiji software (Movie S1, Movie S2, and Movie S3). For whole embryos the mean intensity of a small box outside the embryo where there was no fluorescence (considered as background) was subtracted from the whole worm intensity (region of interest; ROI) using the subtraction function to obtain a background-subtracted image. A polygon tool was used to trace the whole worm (ROI) using the default line width. The mean intensities of whole worms were determined, and background-subtracted image values were plotted.

### Pin enhancer screen

Ahringer library RNAi clones (Geneservice Library) corresponding to the genes listed in File S4 were tested on strain WY964 [*lin-35(n745); sym-4(mn619)*] using standard methods (Kamath and Ahringer, 2003). The RNAi hypersensitive mutation *lin-35(n745)* was used to increase the detection of RNAi enhancers (Lehner et al., 2006; Wang et al., 2005). Control/Empty Vector RNAi-feeding assays were carried out using a bacterial strain carrying the RNAi vector pDF129.36, which produces an ∼200-bp dsRNA not homologous to any *C. elegans* gene (Timmons et al., 2001). Screening was carried out by placing ∼5–10 fourth larval stage (L4) worms (the P0 generation) of strain WY964 onto RNAi plates. The P0s were allowed to develop and have progeny for ∼48 hours, at which point they were removed from the plates. The resulting semi-synchronized F1s were then screened after an additional ∼24 hours using a dissecting microscope to identify plates with an increased frequency of larvae arrested at the first larval stage (L1) with an abnormal (misshapen or bulbous) anterior. After the initial screening, phenotypes for RNAi clones were confirmed a minimum of two times by quantifying the percentage of Pin F1s using a compound microscope. All positive RNAi clones were also sequence confirmed.

### Statistical tests

Statistical tests were performed as outlined (Fay and Gerow, 2013) using GraphPad (Prism). Comparisons between the means of two samples were carried out using the (non-parametric) two-tailed Mann-Whitney test (Figs 3B, 3D, 3E, 4C). In addition, for comparisons of >2 samples, a simple one-way ANOVA was also carried out with results as follows: 3B–Whole embryo (WT, *sym-3*, *sym-4*; P = 0.34); 3B–Anterior perimeter (WT, *sym-3*, *sym-4*; P = 0.027); 3D–Whole embryo (WT, *sym-3*, *sym-4*, *sym-3 sym-4*; P = 0.26); 3D–Anterior perimeter (WT, *sym-3*, *sym-4*, *sym-3 sym-4*; P = 0.72); 3E–FBN-1 (WT, *sym-3*, *sym-4*, *sym-3 sym-4*; P = 0.004); 3E–NOAH-1 (WT, *sym-3*, *sym-4*, *sym-3 sym-4*; P = 0.001). For proportions (Fig 5A) 95% CIs were calculated using the Wilson procedure with a correction for continuity (http://vassarstats.net/prop1.html). For comparisons between two proportions a Fisher’s exact test was used.

## ACKNOWLEDGMENTS

We thank Tony Hyman for recombineered fosmid constructs, Meera Sundaram for strains, Pushph Khanal and John Yochem for help with *mir-51–56* strain constructions, Jimmy Eng and Priska von Haller at the University of Washington Proteomics Resource for their kind help, and Amy Fluet for editing this manuscript.

## SUPPLEMENTAL FIGURE LEGENDS

**File S1. Data for figure panels.**

**File S2. FAM102A and WDR44 LC/MS unfiltered and filtered protein data.**

**File S3. FAM102A and WDR44 LC/MS unfiltered peptide reads.**

**File S4. Enhancer of Pin RNAi screen data.**

**Movie S1. z-Stack movie of FBN-1::sfGFP in a bean-stage embryo.**

**Movie S2. z-Stack movie of FBN-1::sfGFP in an early comma-stage embryo. Movie S3. z-Stack movie of FBN-1::sfGFP in a late comma-stage embryo.**

## SUPPLEMENTAL MATERIALS

**Fig S1.**
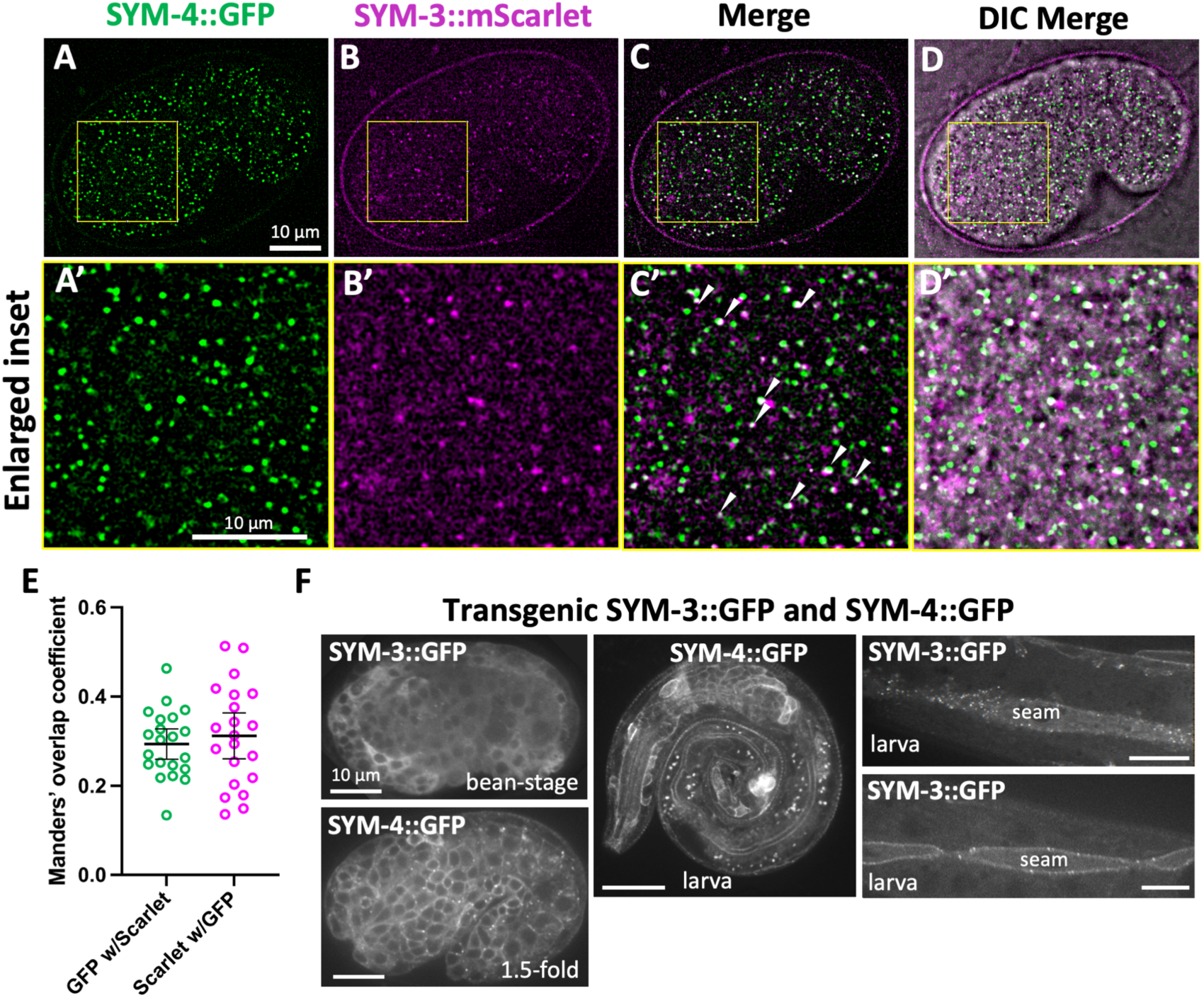
Supplementary images of SYM-3 and SYM-4 expression. (A–D) Expression and colocalization of endogenously tagged SYM-3::mScarlet and SYM-4::GFP in early comma-stage embryos. Anterior is to the left, ventral is down. Arrowheads indicate puncta that overlap (white dots). (A’–D’) Enlarged regions from (A–D) as indicated by the yellow boxes. (E) Manders’ overlap coefficient for SYM-3::mScarlet and SYM-4::GFP embryos. (F) Images of bean-stage and 1.5-fold embryos and larvae expressing SYM-3::GFP and SYM-4::GFP recombineered fosmids from extrachromosomal arrays. Note broad expression and cytoplasmic localization of SYM-3 and SYM-4 including some puncta in SYM-4::GFP embryos and SYM-3::GFP seam cells.

**Fig S2.**
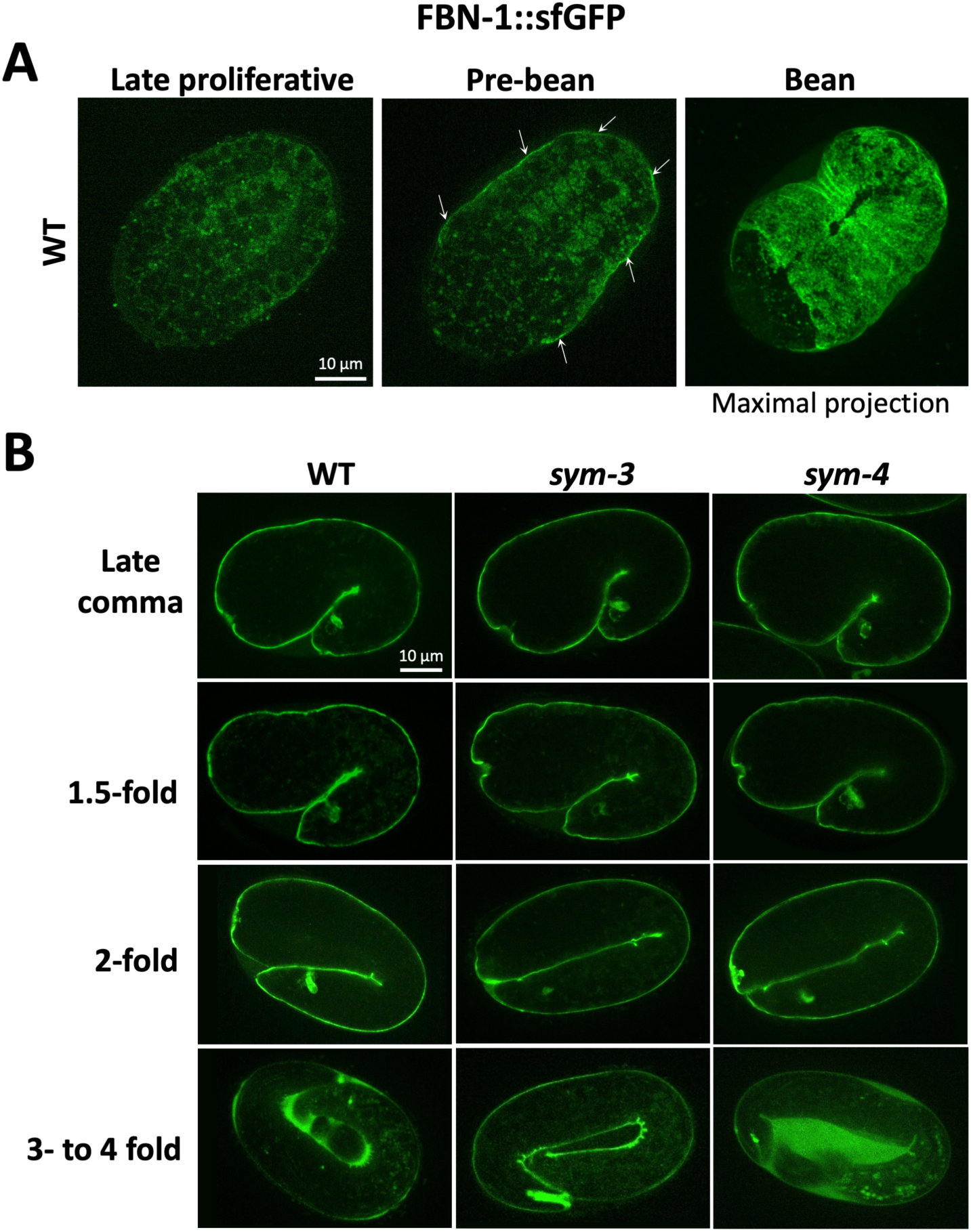
Embryonic expression of FBN-1::sfGFP. (A) Endogenously tagged FBN-1::sfGFP in representative wild-type (WT) embryos at the indicated stages. Anterior is to the left, ventral is down. Arrows (middle panel) indicate the appearance of apical GFP, coincident with the onset of morphogenesis. Note FBN-1::sfGFP beginning to cover the apical surface in the maximal-projection image of the bean-stage embryo (rightmost panel) The absence of FBN-1::sfGFP signal in the anterior is consistent with a lack of anterior epidermal cells at this stage; epidermal cells complete migration to and containment of the anterior by the comma stage (Grimbert et al., 2021). (B) Representative FBN-1::sfGFP expression in wild-type, *sym-3*, and *sym-4* embryos at the indicated stages.

**Fig S3.**
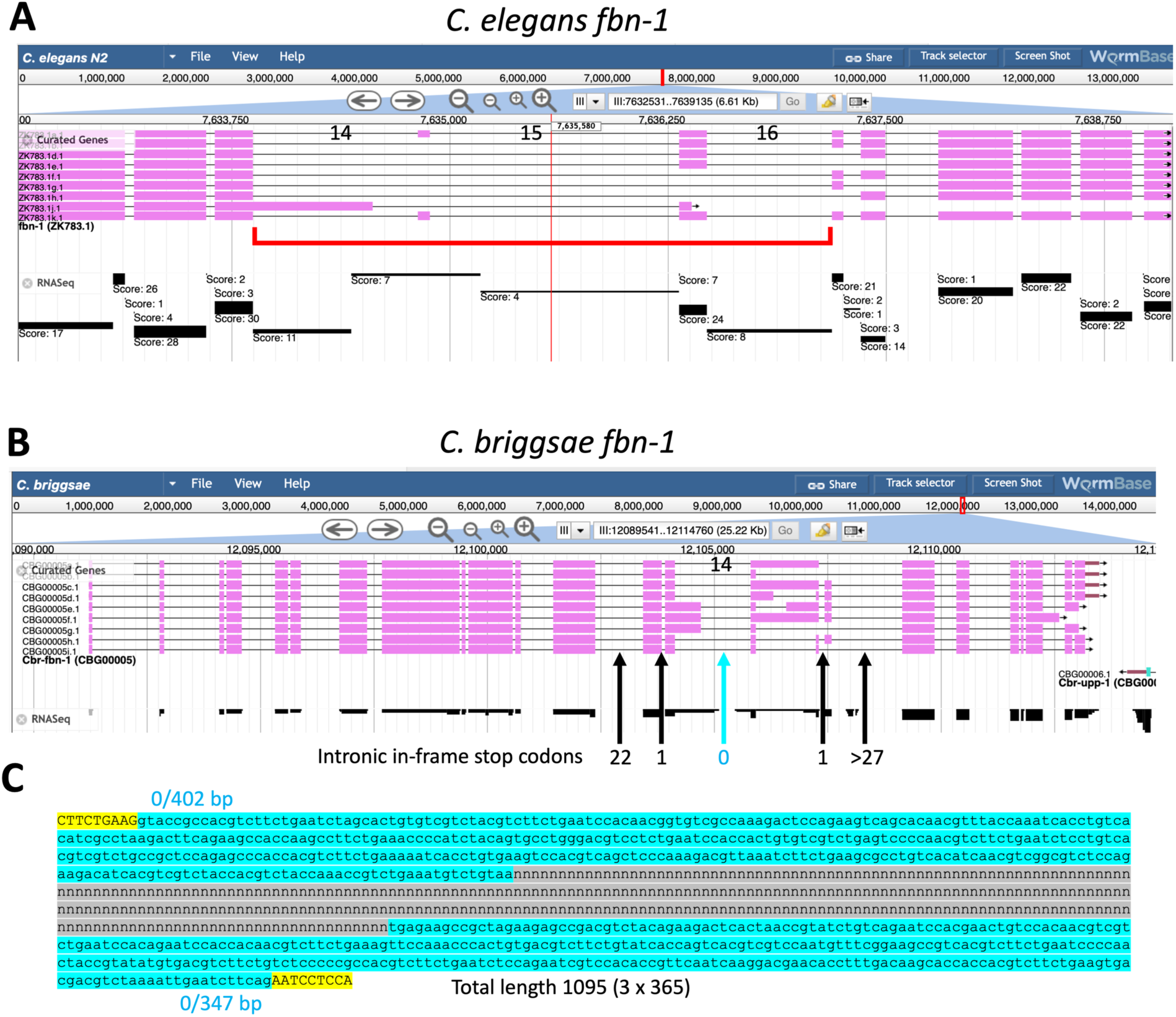
Genomic loci of *C. elegans* and *C. briggsae fbn-1*. (A, B) Images taken from WormBase of the *C. elegans* (A) and *C. briggsae* (B) *fbn-1* genomic loci. RNAseq data are displayed below the pink gene diagrams (black lines and bars). RNAseq scores correspond to read numbers. The *C. elegans fbn-1* locus is enlarged to highlight the relevant region (red bracket), with introns 14–16 indicated. For *C. briggsae fbn-1*, the number of intronic in-frame stop codons is indicated (arrows). Note the absence of detected in-frame stop codons in intron 14 (blue arrow).(C) Available sequence on WormBase for intron 14 of *C. briggsae fbn-1*, which includes an unassigned 346-bp central region (gray background). The regions downstream of exon 14 (402 bp) and upstream of exon 15 (347 bp) form a continuous open reading frame (blue background). Yellow background indicates exons 14 and 15.

## REFERENCES

**Bhuin**, **T.** **and** **Roy**, **J. K**. (2015). Rab11 in disease progression. Int J Mol Cell Med 4, 1–8.

**Casamento**, **A.** **and** **Boucrot**, **E**. (2020). Molecular mechanism of Fast Endophilin-Mediated Endocytosis. Biochem J 477, 2327–2345.

**Ceron**, **J.**, **Rual**, **J. F.**, **Chandra**, **A.**, **Dupuy**, **D.**, **Vidal**, **M.** **and** **van den Heuvel**, **S.** (2007). Large-scale RNAi screens identify novel genes that interact with the C. elegans retinoblastoma pathway as well as splicing-related components with synMuv B activity. BMC Dev Biol 7, 30.

**Chisholm**, **A. D.** **and** **Hardin**, **J**. (2005). Epidermal morphogenesis. WormBook, 1-22.

**Chisholm**, **A. D.** **and** **Hsiao**, **T. I**. (2012). The Caenorhabditis elegans epidermis as a model skin. I: development, patterning, and growth. Wiley Interdiscip Rev Dev Biol 1, 861–878.

**Choi**, **M. K.**, **Son**, **S.**, **Hong**, **M.**, **Choi**, **M. S.**, **Kwon**, **J. Y.** **and** **Lee**, **J**. (2016). Maintenance of Membrane Integrity and Permeability Depends on a Patched-Related Protein in Caenorhabditis elegans. Genetics 202, 1411–1420.

**Chung**, **Y. C.**, **Wei**, **W. C.**, **Huang**, **S. H.**, **Shih**, **C. M.**, **Hsu**, **C. P.**, **Chang**, **K. J.** **and** **Chao**, **W. T**. (2014). Rab11 regulates E-cadherin expression and induces cell transformation in colorectal carcinoma. BMC Cancer 14, 587.

**Cohen**, **J. D.**, **Bermudez**, **J. G.**, **Good**, **M. C.** **and** **Sundaram**, **M. V**. (2020a). A C. elegans Zona Pellucida domain protein functions via its ZPc domain. PLoS Genet 16, e1009188.

**Cohen**, **J. D.**, **Flatt**, **K. M.**, **Schroeder**, **N. E.** **and** **Sundaram**, **M. V**. (2019). Epithelial Shaping by Diverse Apical Extracellular Matrices Requires the Nidogen Domain Protein DEX-1 in Caenorhabditis elegans. Genetics 211, 185–200.

**Cohen**, **J. D.**, **Sparacio**, **A. P.**, **Belfi**, **A. C.**, **Forman-Rubinsky**, **R.**, **Hall**, **D. H.**, **Maul-Newby**, **H.**, **Frand**, **A. R.** **and** **Sundaram**, **M. V**. (2020b). A multi-layered and dynamic apical extracellular matrix shapes the vulva lumen in Caenorhabditis elegans. Elife 9.

**Cohen**, **J. D.** **and** **Sundaram**, **M. V**. (2020). C. elegans Apical Extracellular Matrices Shape Epithelia. J Dev Biol 8, 1–26.

**Davies**, **A. G.**, **Spike**, **C. A.**, **Shaw**, **J. E.** **and** **Herman**, **R. K**. (1999). Functional overlap between the mec-8 gene and five sym genes in Caenorhabditis elegans. Genetics 153, 117–134.

**Fan**, **X.**, **She**, **Y. M.**, **Bagshaw**, **R. D.**, **Callahan**, **J. W.**, **Schachter**, **H.** **and** **Mahuran**, **D. J**. (2005). Identification of the hydrophobic glycoproteins of Caenorhabditis elegans. Glycobiology 15, 952–964.

**Fay**, **D. S.** **and** **Gerow**, **K**. (2013). A biologist’s guide to statistical thinking and analysis. WormBook, 1-54.

**Fay**, **D. S.**, **Qiu**, **X.**, **Large**, **E.**, **Smith**, **C. P.**, **Mango**, **S.** **and** **Johanson**, **B. L**. (2004). The coordinate regulation of pharyngeal development in C. elegans by lin-35/Rb, pha-1, and ubc-18. Dev Biol 271, 11–25.

**Ferro**, **E.**, **Bosia**, **C.** **and** **Campa**, **C. C**. (2021). RAB11-Mediated Trafficking and Human Cancers: An Updated Review. Biology (Basel) 10.

**Forman-Rubinsky**, **R.**, **Cohen**, **J. D.** **and** **Sundaram**, **M. V**. (2017). Lipocalins Are Required for Apical Extracellular Matrix Organization and Remodeling in Caenorhabditis elegans. Genetics 207, 625–642.

**Frand**, **A. R.**, **Russel**, **S.** **and** **Ruvkun**, **G**. (2005). Functional genomic analysis of C. elegans molting. PLoS Biol 3, e312.

**Gibieza**, **P.** **and** **Petrikaite**, **V**. (2021). The dual functions of Rab11 and Rab35 GTPases-regulation of cell division and promotion of tumorigenicity. Am J Cancer Res 11, 1861–1872.

**Gillon**, **A. D.**, **Latham**, **C. F.** **and** **Miller**, **E. A**. (2012). Vesicle-mediated ER export of proteins and lipids. Biochim Biophys Acta 1821, 1040–1049.

**Goldenring**, **J. R**. (2019). Membrane Trafficking Decisions Regulate Primary Cilium Formation. Trends Cell Biol 29, 607–608.

**Goldstein**, **B.** **and** **Nance**, **J**. (2020). Caenorhabditis elegans Gastrulation: A Model for Understanding How Cells Polarize, Change Shape, and Journey Toward the Center of an Embryo. Genetics 214, 265–277.

**Granato**, **M.**, **Schnabel**, **H.** **and** **Schnabel**, **R**. (1994). pha-1, a selectable marker for gene transfer in C. elegans. Nucleic Acids Res 22, 1762–1763.

**Griffin**, **C. T.**, **Trejo**, **J.** **and** **Magnuson**, **T**. (2005). Genetic evidence for a mammalian retromer complex containing sorting nexins 1 and 2. Proc Natl Acad Sci U S A 102, 15173–15177.

**Grimbert**, **S.**, **Mastronardi**, **K.**, **Richard**, **V.**, **Christensen**, **R.**, **Law**, **C.**, **Zardoui**, **K.**, **Fay**, **D.** **and** **Piekny**, **A**. (2021). Multi-tissue patterning drives anterior morphogenesis of the C. elegans embryo. Dev Biol 471, 49–64.

**Guichard**, **A.**, **Nizet**, **V.** **and** **Bier**, **E**. (2014). RAB11-mediated trafficking in host-pathogen interactions. Nat Rev Microbiol 12, 624–634.

**Gullapalli**, **A.**, **Garrett**, **T. A.**, **Paing**, **M. M.**, **Griffin**, **C. T.**, **Yang**, **Y.** **and** **Trejo**, **J**. (2004). A role for sorting nexin 2 in epidermal growth factor receptor down-regulation: evidence for distinct functions of sorting nexin 1 and 2 in protein trafficking. Mol Biol Cell 15, 2143–2155.

**Heiman**, **M. G.** **and** **Shaham**, **S**. (2009). DEX-1 and DYF-7 establish sensory dendrite length by anchoring dendritic tips during cell migration. Cell 137, 344–355.

**Hendriks**, **G. J.**, **Gaidatzis**, **D.**, **Aeschimann**, **F.** **and** **Grosshans**, **H**. (2014). Extensive oscillatory gene expression during C. elegans larval development. Mol Cell 53, 380–392.

**Hernandez**, **M. R.** **and** **Ye**, **H**. (1993). Glaucoma: changes in extracellular matrix in the optic nerve head. Ann Med 25, 309–315.

**Johnstone**, **I. L**. (1994). The cuticle of the nematode Caenorhabditis elegans: a complex collagen structure. Bioessays 16, 171–178.

**Jones**, **M. R.**, **Rose**, **A. M.** **and** **Baillie**, **D. L**. (2013). The ortholog of the human proto-oncogene ROS1 is required for epithelial development in C. elegans. Genesis 51, 545–561.

**Kaji**, **H.**, **Kamiie**, **J.**, **Kawakami**, **H.**, **Kido**, **K.**, **Yamauchi**, **Y.**, **Shinkawa**, **T.**, **Taoka**, **M.**, **Takahashi**, **N.** **and** **Isobe**, **T**. (2007). Proteomics reveals N-linked glycoprotein diversity in Caenorhabditis elegans and suggests an atypical translocation mechanism for integral membrane proteins. Mol Cell Proteomics 6, 2100–2109.

**Kamath**, **R. S.** **and** **Ahringer**, **J**. (2003). Genome-wide RNAi screening in Caenorhabditis elegans. Methods 30, 313–321.

**Kang**, **S. H.** **and** **Kramer**, **J. M**. (2000). Nidogen is nonessential and not required for normal type IV collagen localization in Caenorhabditis elegans. Mol Biol Cell 11, 3911–3923.

**Katz**, **S. S.**, **Barker**, **T. J.**, **Maul-Newby**, **H. M.**, **Sparacio**, **A. P.**, **Nguyen**, **K. C. Q.**, **Maybrun**, **C. L.**, **Belfi**, **A.**, **Cohen**, **J. D.**, **Hall**, **D. H.**, **Sundaram**, **M. V.**, et al. (2022). A transient apical extracellular matrix relays cytoskeletal patterns to shape permanent acellular ridges on the surface of adult C. elegans. PLoS Genet 18, e1010348.

**Kelley**, **M.**, **Yochem**, **J.**, **Krieg**, **M.**, **Calixto**, **A.**, **Heiman**, **M. G.**, **Kuzmanov**, **A.**, **Meli**, **V.**, **Chalfie**, **M.**, **Goodman**, **M. B.**, **Shaham**, **S.**, et al. (2015). FBN-1, a fibrillin-related protein, is required for resistance of the epidermis to mechanical deformation during C. elegans embryogenesis. Elife 4.

**Khor**, **C. C.**, **Do**, **T.**, **Jia**, **H.**, **Nakano**, **M.**, **George**, **R.**, **Abu-Amero**, **K.**, **Duvesh**, **R.**, **Chen**, **L. J.**, **Li**, **Z.**, **Nongpiur**, **M. E.**, et al. (2016). Genome-wide association study identifies five new susceptibility loci for primary angle closure glaucoma. Nat Genet 48, 556–562.

**Kjaerulff**, **O.**, **Brodin**, **L.** **and** **Jung**, **A**. (2011). The structure and function of endophilin proteins. Cell Biochem Biophys 60, 137–154.

**Kuzmanov**, **A.**, **Yochem**, **J.** **and** **Fay**, **D. S**. (2014). Analysis of PHA-1 reveals a limited role in pharyngeal development and novel functions in other tissues. Genetics 198, 259–268.

**Labouesse**, **M**. (2012). Role of the extracellular matrix in epithelial morphogenesis: a view from C. elegans. Organogenesis 8, 65–70.

**Lamkin**, **E. R.** **and** **Heiman**, **M. G**. (2017). Coordinated morphogenesis of neurons and glia. Curr Opin Neurobiol 47, 58–64.

**Lehner**, **B.**, **Calixto**, **A.**, **Crombie**, **C.**, **Tischler**, **J.**, **Fortunato**, **A.**, **Chalfie**, **M.** **and** **Fraser**, **A. G**. (2006). Loss of LIN-35, the Caenorhabditis elegans ortholog of the tumor suppressor p105Rb, results in enhanced RNA interference. Genome Biol 7, R4.

**Li**, **K.**, **Yang**, **C.**, **Wan**, **X.**, **Xu**, **J.**, **Luo**, **Q.**, **Cheng**, **Y.**, **Peng**, **J.**, **Gong**, **B.**, **Jiang**, **L.**, **Liu**, **Y.**, **et al**. (2020). Evaluation of the association between five genetic variants and primary open-angle glaucoma in a Han Chinese population. Ophthalmic Genet 41, 252–256.

**Li** **Zheng**, **S.**, **Adams**, **J. G.** **and Chisholm**, **A.** **D**. (2020). Form and function of the apical extracellular matrix: new insights from Caenorhabditis elegans, Drosophila melanogaster, and the vertebrate inner ear. Fac Rev 9, 27.

**Longman**, **D.**, **Arrisi**, **P.**, **Johnstone**, **I. L.** **and** **Caceres**, **J. F**. (2008). Chapter 7. Nonsense-mediated mRNA decay in Caenorhabditis elegans. Methods Enzymol 449, 149–164.

**Lord**, **C.**, **Ferro-Novick**, **S.** **and** **Miller**, **E. A**. (2013). The highly conserved COPII coat complex sorts cargo from the endoplasmic reticulum and targets it to the golgi. Cold Spring Harb Perspect Biol 5.

**Lucken-Ardjomande Hasler**, **S.**, **Vallis**, **Y.**, **Pasche**, **M.** and **McMahon**, **H.** **T**. (2020). GRAF2, WDR44, and MICAL1 mediate Rab8/10/11-dependent export of E-cadherin, MMP14, and CFTR DeltaF508. J Cell Biol 219.

**Lundquist**, **E. A.**, **Herman**, **R. K.**, **Rogalski**, **T. M.**, **Mullen**, **G. P.**, **Moerman**, **D. G.** **and** **Shaw**, **J. E**. (1996). The mec-8 gene of C. elegans encodes a protein with two RNA recognition motifs and regulates alternative splicing of unc-52 transcripts. Development 122, 1601–1610.

**Lykke-Andersen**, **S.** **and** **Jensen**, **T. H**. (2015). Nonsense-mediated mRNA decay: an intricate machinery that shapes transcriptomes. Nat Rev Mol Cell Biol 16, 665–677.

**Lynch**, **K. L.**, **Gerona**, **R. R.**, **Larsen**, **E. C.**, **Marcia**, **R. F.**, **Mitchell**, **J. C.** **and** **Martin**, **T. F**. (2007). Synaptotagmin C2A loop 2 mediates Ca2+-dependent SNARE interactions essential for Ca2+-triggered vesicle exocytosis. Mol Biol Cell 18, 4957–4968.

**Maduro**, **M.** **and** **Pilgrim**, **D**. (1996). Conservation of function and expression of unc-119 from two Caenorhabditis species despite divergence of non-coding DNA. Gene 183, 77–85.

**Mammoto**, **A.**, **Ohtsuka**, **T.**, **Hotta**, **I.**, **Sasaki**, **T.** **and** **Takai**, **Y**. (1999). Rab11BP/Rabphilin-11, a downstream target of rab11 small G protein implicated in vesicle recycling. J Biol Chem 274, 25517–25524.

**Mammoto**, **A.**, **Sasaki**, **T.**, **Kim**, **Y.** **and** **Takai**, **Y**. (2000). Physical and functional interaction of rabphilin-11 with mammalian Sec13 protein. Implication in vesicle trafficking. J Biol Chem 275, 13167–13170.

**Meeuse**, **M. W.**, **Hauser**, **Y. P.**, **Morales Moya**, **L. J.**, **Hendriks**, **G. J.**, **Eglinger**, **J.**, **Bogaarts**, **G.**, **Tsiairis**, **C.** **and** **Grosshans**, **H**. (2020). Developmental function and state transitions of a gene expression oscillator in Caenorhabditis elegans. Mol Syst Biol 16, e9498.

**Mello**, **C. C.**, **Kramer**, **J. M.**, **Stinchcomb**, **D.** **and** **Ambros**, **V**. (1991). Efficient gene transfer in C.elegans: extrachromosomal maintenance and integration of transforming sequences. EMBO J 10, 3959–3970.

**Muir**, **V. S.**, **Gasch**, **A. P.** **and** **Anderson**, **P**. (2018). The Substrates of Nonsense-Mediated mRNA Decay in Caenorhabditis elegans. G3(Bethesda) 8, 195-205.

**Nikhalashree**, **S.**, **George**, **R.**, **Shantha**, **B.**, **Lingam**, **V.**, **Vidya**, **W.**, **Panday**, **M.**, **Sulochana**, **K. N.** **and** **Coral**, **K**. (2019). Detection of Proteins Associated with Extracellular Matrix Regulation in the Aqueous Humour of Patients with Primary Glaucoma. Curr Eye Res 44, 1018–1025.

**Nongpiur**, **M. E.**, **Cheng**, **C. Y.**, **Duvesh**, **R.**, **Vijayan**, **S.**, **Baskaran**, **M.**, **Khor**, **C. C.**, **Allen**, **J.**, **Kavitha**, **S.**, **Venkatesh**, **R.**, **Goh**, **D.**, et al. (2018). Evaluation of Primary Angle-Closure Glaucoma Susceptibility Loci in Patients with Early Stages of Angle-Closure Disease. Ophthalmology 125, 664–670.

**O’Callaghan**, **J.**, **Cassidy**, **P. S.** **and** **Humphries**, **P**. (2017). Open-angle glaucoma: therapeutically targeting the extracellular matrix of the conventional outflow pathway. Expert Opin Ther Targets 21, 1037–1050.

**Olsen**, **J. V.** **and** **Mann**, **M**. (2013). Status of large-scale analysis of post-translational modifications by mass spectrometry. Mol Cell Proteomics 12, 3444–3452.

**Page**, **A. P.** **and** **Johnstone**, **I. L**. (2007). The cuticle. WormBook, 1-15.

**Portereiko**, **M. F.** **and** **Mango**, **S. E**. (2001). Early morphogenesis of the Caenorhabditis elegans pharynx. Dev Biol 233, 482–494.

**Priess**, **J. R.** **and** **Hirsh**, **D. I**. (1986). Caenorhabditis elegans morphogenesis: the role of the cytoskeleton in elongation of the embryo. Dev Biol 117, 156–173.

**Ramel**, **D.**, **Wang**, **X.**, **Laflamme**, **C.**, **Montell**, **D. J.** **and** **Emery**, **G**. (2013). Rab11 regulates cell-cell communication during collective cell movements. Nat Cell Biol 15, 317–324.

**Roberg**, **K. J.**, **Rowley**, **N.** **and** **Kaiser**, **C. A**. (1997). Physiological regulation of membrane protein sorting late in the secretory pathway of Saccharomyces cerevisiae. J Cell Biol 137, 1469–1482.

**Saleh**, **M. C.**, **van Rij**, **R. P.**, **Hekele**, **A.**, **Gillis**, **A.**, **Foley**, **E.**, **O’Farrell**, **P. H.** **and** **Andino**, **R.** (2006). The endocytic pathway mediates cell entry of dsRNA to induce RNAi silencing. Nat Cell Biol 8, 793–802.

**Sapio**, **M. R.**, **Hilliard**, **M. A.**, **Cermola**, **M.**, **Favre**, **R.** **and** **Bazzicalupo**, **P**. (2005). The Zona Pellucida domain containing proteins, CUT-1, CUT-3 and CUT-5, play essential roles in the development of the larval alae in Caenorhabditis elegans. Dev Biol 282, 231-245.

**Sarov**, **M.**, **Murray**, **J. I.**, **Schanze**, **K.**, **Pozniakovski**, **A.**, **Niu**, **W.**, **Angermann**, **K.**, **Hasse**, **S.**, **Rupprecht**, **M.**, **Vinis**, **E.**, **Tinney**, **M.**, et al. (2012). A genome-scale resource for in vivo tag-based protein function exploration in C. elegans. Cell 150, 855–866.

**Seaman**, **M. N**. (2012). The retromer complex - endosomal protein recycling and beyond. J Cell Sci 125, 4693–4702.

---- (2021). The Retromer Complex: From Genesis to Revelations. Trends Biochem Sci 46, 608–620.

**Sharma**, **V.**, **Eng**, **J. K.**, **Maccoss**, **M. J.** **and** **Riffle**, **M**. (2012). A mass spectrometry proteomics data management platform. Mol Cell Proteomics 11, 824–831.

**Shaw**, **W. R.**, **Armisen**, **J.**, **Lehrbach**, **N. J.** **and** **Miska**, **E. A**. (2010). The conserved miR-51 microRNA family is redundantly required for embryonic development and pharynx attachment in Caenorhabditis elegans. Genetics 185, 897–905.

**Shi**, **A.**, **Chen**, **C. C.**, **Banerjee**, **R.**, **Glodowski**, **D.**, **Audhya**, **A.**, **Rongo**, **C.** **and** **Grant**, **B. D**. (2010). EHBP-1 functions with RAB-10 during endocytic recycling in Caenorhabditis elegans. Mol Biol Cell 21, 2930–2943.

**Shi**, **H.**, **Chen**, **Y.**, **Lu**, **H.**, **Zhu**, **R.**, **Zhang**, **J.**, **He**, **M.** **and** **Guan**, **H**. (2021). In-depth analysis of eight susceptibility loci of primary angle closure glaucoma in Han Chinese. Exp Eye Res 202, 108350.

**Simmer**, **F.**, **Moorman**, **C.**, **van der Linden**, **A. M.**, **Kuijk**, **E**., van den Berghe, P. V., Kamath, R. S., Fraser, A. G., Ahringer, J. and Plasterk, R. H. (2003). Genome-wide RNAi of C. elegans using the hypersensitive rrf-3 strain reveals novel gene functions. PLoS Biol 1, E12.

**Spanier**, **B.**, **Sturzenbaum**, **S. R.**, **Holden-Dye**, **L. M.** **and** **Baumeister**, **R**. (2005). Caenorhabditis elegans neprilysin NEP-1: an effector of locomotion and pharyngeal pumping. J Mol Biol 352, 429–437.

**Spike**, **C. A.**, **Davies**, **A. G.**, **Shaw**, **J. E.** **and** **Herman**, **R. K**. (2002). MEC-8 regulates alternative splicing of unc-52 transcripts in C. elegans hypodermal cells. Development 129, 4999–5008.

**Stein**, **L.**, **Sternberg**, **P.**, **Durbin**, **R.**, **Thierry-Mieg**, **J.** **and** **Spieth**, **J**. (2001). WormBase: network access to the genome and biology of Caenorhabditis elegans. Nucleic Acids Res 29, 82–86.

**Suh**, **J.** **and** **Hutter**, **H**. (2012). A survey of putative secreted and transmembrane proteins encoded in the C. elegans genome. BMC Genomics 13, 333.

**Sundaram**, **M. V.** **and** **Cohen**, **J. D**. (2017). Time to make the doughnuts: Building and shaping seamless tubes. Semin Cell Dev Biol 67, 123–131.

**Thein**, **M. C.**, **Winter**, **A. D.**, **Stepek**, **G.**, **McCormack**, **G.**, **Stapleton**, **G.**, **Johnstone**, **I. L.** **and** **Page**, **A. P**. (2009). Combined extracellular matrix cross-linking activity of the peroxidase MLT-7 and the dual oxidase BLI-3 is critical for post-embryonic viability in Caenorhabditis elegans. J Biol Chem 284, 17549–17563.

**Thibodeau**, **M. C.**, **Harris**, **N. J.**, **Jenkins**, **M. L.**, **Parson**, **M. A. H.**, **Evans**, **J. T.**, **Scott**, **M. K.**, **Shaw**, **A. L.**, **Pokorny**, **D.**, **Leonard**, **T. A.** **and** **Burke**, **J. E**. (2023). Molecular basis for the recruitment of the Rab effector protein WDR44 by the GTPase Rab11. J Biol Chem 299, 102764.

**Timmons**, **L.**, **Court**, **D. L.** **and** **Fire**, **A**. (2001). Ingestion of bacterially expressed dsRNAs can produce specific and potent genetic interference in Caenorhabditis elegans. Gene 263, 103–112.

**Topalidou**, **I.**, **Cattin-Ortola**, **J.**, **Pappas**, **A. L.**, **Cooper**, **K.**, **Merrihew**, **G. E.**, **MacCoss**, **M. J.** **and** **Ailion**, **M**. (2016). The EARP Complex and Its Interactor EIPR-1 Are Required for Cargo Sorting to Dense-Core Vesicles. PLoS Genet 12, e1006074.

**Tribble**, **J. R.**, **Williams**, **P. A.**, **Caterson**, **B.**, **Sengpiel**, **F.** **and** **Morgan**, **J. E**. (2018). Digestion of the glycosaminoglycan extracellular matrix by chondroitinase ABC supports retinal ganglion cell dendritic preservation in a rodent model of experimental glaucoma. Mol Brain 11, 69.

**Tu**, **Y.** **and** **Seaman**, **M. N. J**. (2021). Navigating the Controversies of Retromer-Mediated Endosomal Protein Sorting. Front Cell Dev Biol 9, 658741.

**Vranka**, **J. A.**, **Kelley**, **M. J.**, **Acott**, **T. S.** **and** **Keller**, **K. E**. (2015). Extracellular matrix in the trabecular meshwork: intraocular pressure regulation and dysregulation in glaucoma. Exp Eye Res 133, 112–125.

**Vuong-Brender**, **T. T.**, **Yang**, **X.** **and** **Labouesse**, **M**. (2016). C. elegans Embryonic Morphogenesis. Curr Top Dev Biol 116, 597–616.

**Vuong-Brender**, **T. T. K.**, **Suman**, **S. K.** **and** **Labouesse**, **M**. (2017). The apical ECM preserves embryonic integrity and distributes mechanical stress during morphogenesis. Development 144, 4336–4349.

**Walia**, **V.**, **Cuenca**, **A.**, **Vetter**, **M.**, **Insinna**, **C.**, **Perera**, **S.**, **Lu**, **Q.**, **Ritt**, **D. A.**, **Semler**, **E.**, **Specht**, **S.**, **Stauffer**, **J**., et al. (2019). Akt Regulates a Rab11-Effector Switch Required for Ciliogenesis. Dev Cell 50, 229–246 e227.

**Wang**, **D.**, **Kennedy**, **S.**, **Conte**, **D.**, Jr., **Kim**, **J. K.**, **Gabel**, **H. W.**, **Kamath**, **R. S.**, **Mello**, **C. C.** **and** **Ruvkun**, **G**. (2005). Somatic misexpression of germline P granules and enhanced RNA interference in retinoblastoma pathway mutants. Nature 436, 593–597.

**Wang**, **P.**, **Liu**, **H.**, **Wang**, **Y.**, **Liu**, **O.**, **Zhang**, **J.**, **Gleason**, **A.**, **Yang**, **Z.**, **Wang**, **H.**, **Shi**, **A.** **and** **Grant**, **B. D**. (2016). RAB-10 Promotes EHBP-1 Bridging of Filamentous Actin and Tubular Recycling Endosomes. PLoS Genet 12, e1006093.

**Welz**, **T.**, **Wellbourne-Wood**, **J.** **and** **Kerkhoff**, **E**. (2014). Orchestration of cell surface proteins by Rab11. Trends Cell Biol 24, 407–415.

**Wiggs**, **J. L**. (2015). Glaucoma Genes and Mechanisms. Prog Mol Biol Transl Sci 134, 315–342.

**Yochem**, **J.**, **Bell**, **L. R.** **and** **Herman**, **R. K**. (2004). The identities of sym-2, sym-3 and sym-4, three genes that are synthetically lethal with mec-8 in Caenorhabditis elegans. Genetics 168, 1293–1306.

**Zeng**, **J.**, **Ren**, **M.**, **Gravotta**, **D.**, **De Lemos-Chiarandini**, **C.**, **Lui**, **M.**, **Erdjument-Bromage**, **H.**, **Tempst**, **P.**, **Xu**, **G.**, **Shen**, **T. H.**, **Morimoto**, **T**., et al. (1999). Identification of a putative effector protein for rab11 that participates in transferrin recycling. Proc Natl Acad Sci U S A 96, 2840–2845.

**Zhang**, **D.** **and** **Aravind**, **L**. (2010). Identification of novel families and classification of the C2 domain superfamily elucidate the origin and evolution of membrane targeting activities in eukaryotes. Gene 469, 18–30.

**Zhang**, **J.**, **Su**, **G.**, **Wu**, **Q.**, **Liu**, **J.**, **Tian**, **Y.**, **Liu**, **X.**, **Zhou**, **J.**, **Gao**, **J.**, **Chen**, **W.**, **Chen**, **D.**, **et al**. (2021). Rab11-mediated recycling endosome role in nervous system development and neurodegenerative diseases. Int J Neurosci 131, 1012–1018.

**Zhuang**, **W.**, **Wang**, **S.**, **Hao**, **J.**, **Xu**, **M.**, **Chi**, **H.**, **Piao**, **S.**, **Ma**, **J.**, **Zhang**, **X.** **and** **Ha**, **S**. (2018). Genotype-ocular biometry correlation analysis of eight primary angle closure glaucoma susceptibility loci in a cohort from Northern China. PLoS One 13, e0206935.

**Zugasti**, **O.**, **Rajan**, **J.** **and** **Kuwabara**, **P. E**. (2005). The function and expansion of the Patched- and Hedgehog-related homologs in C. elegans. Genome Res 15, 1402–1410.

